# Fibronectin-integrin α5 signaling promotes thoracic aortic aneurysm in a mouse model of Marfan syndrome

**DOI:** 10.1101/2022.08.16.504169

**Authors:** Minghao Chen, Cristina Cavinato, Jens Hansen, Keiichiro Tanaka, Pengwei Ren, Abdulrahman Hassab, David S. Li, Eric. Joshuao, George Tellides, Ravi Iyengar, Jay D. Humphrey, Martin A. Schwartz

**Affiliations:** Cardiovascular Research Center, Yale School of Medicine, New Haven, CT, USA; Department of Biomedical Engineering, Yale University, New Haven, CT, USA; Department of Pharmacological Sciences and Institute for Systems Biomedicine, Icahn School of Medicine at Mount Sinai, New York, NY, 10029, USA; Department of Surgery, Yale School of Medicine, New Haven, CT, USA; Vascular Biology and Therapeutics Program, Yale School of Medicine, New Haven, CT, USA; Departments of Medicine (Cardiology) and Cell Biology, Yale School of Medicine, New Haven, CT, USA

**Keywords:** thoracic aneurysm, Marfan syndrome, fibronectin, integrin α5, smooth muscle cell, inflammation, NF-kB

## Abstract

**Background:** Marfan syndrome, caused by mutations in the gene for the extracellular matrix (ECM) glycoprotein fibrillin-1, leads to thoracic aortic aneurysms (TAAs). Phenotypic modulation of vascular smooth muscle cells (SMCs) and ECM remodeling are characteristics of both non-syndromic and Marfan aneurysms. The ECM protein fibronectin (FN) is elevated in the tunica media of TAAs and amplifies inflammatory signaling in endothelial and SMCs through its main receptor, integrin α5β1. We investigated the role of integrin α5-specific signals in Marfan mice in which the cytoplasmic domain of integrin α5 was replaced with that of integrin α2 (denoted α5/2 chimera).

**Methods:** We used α5/2 chimera mouse crossed with *Fbn1^mgR/mgR^* genetic background (mgR, a mouse model of Marfan syndrome) to compare the survival rate and pathogenesis of TAAs among wild type, α5/2, mgR and α5/2; mgR mice. Further biochemical and microscopic analysis of porcine and mouse aortic SMCs allowed us to identify the molecular mechanisms by which FN affects SMCs and subsequent development of TAAs.

**Results:** FN was elevated in the thoracic aortas from Marfan patients, in non-syndromic aneurysms and in the mgR mouse model of Marfan syndrome. The α5/2 mutation greatly prolonged survival of Marfan mice, with improved elastic fiber integrity, mechanical properties, SMC density, and SMC contractile gene expression. Furthermore, *in vitro,* plating of wild-type, but not α5/2, SMCs on FN decreased contractile gene expression and activated inflammatory pathways. These effects correlated with increased NF-kB activation and immune cell infiltration in the mgR aortas, which was rescued in the α5/2 mgR aortas.

**Conclusions:** FN-integrin α5 signaling is a significant driver of TAA in the mgR mouse model. This pathway warrants further investigation as a therapeutic target.

## INTRODUCTION

Approximately 1 in 5000 people are afflicted with Marfan syndrome (MFS), a genetic disorder that affects connective tissues throughout the body. Despite improved patient outcomes due to advances in medical genetics, medical imaging, and prophylactic surgery, MFS leads to aneurysms of the aortic root and ascending aorta that can dissect or rupture, resulting in significant morbidity and mortality. MFS results from mutations in the *FBN1* gene that encodes the extracellular matrix glycoprotein fibrillin-1, which normally associates with elastin to form stable elastic fibers. Fibrillin-1 also regulates transforming growth factor-beta (TGFβ) bioavailability (1) while fibrillin-1 microfibers form a key connection between the elastic laminae and medial smooth muscle cells (SMCs) to facilitate mechanosensing by these cells (2). Dysfunctional mechanosensing appears to be a significant driver of the progressive deterioration of aortic composition and mechanical properties that ultimately results in aneurysmal dilatation or catastrophic failure (3–5).

Although it has been 30+ years since the discovery of the monogenic cause of MFS (6), recent bulk and single cell sequencing reveal hundreds of differentially expressed genes in the two most common mouse models of MFS, a hypomorphic *Fbn1^mgR/mgR^* mouse (7) and a less severe knock-in *Fbn1^C1041G/+^* mouse (8). Gene ontology studies highlight, among other processes affected, altered cell adhesion, extracellular matrix-receptor interactions, and focal adhesions. Amongst the many increased transcripts are those for FN (*Fn1*) and the β1 integrin subunit (*Itgb1*), which pairs with the α5 subunit to form the high affinity FN receptor integrin α5β1 (9). α5β1 also binds fibrillin-1, as do multiple αv integrins (10). Mice deficient in the focal adhesion protein integrin linked kinase, which is implicated in mechanosensing, develop thoracic aortic aneurysms (TAAs) (11), while a recent genome wide screen of over 500 patients with sporadic TAA identified associations with additional focal adhesion genes, enforcing the link between integrin-mediated mechanosensing and aneurysms (12).

Fibronectin plays multiple roles within the extracellular matrix, including initiating fibrillin-1 assembly (13), promoting collagen fibril assembly (14, 15), and, along with fibrillin-1, sequestering latent TGFβ (16). Conversely, TGFβ increases fibronectin deposition by many cell types including SMCs (17, 18). Increased staining for FN was detected in human TAAs (19). Multiple studies have demonstrated that FN promotes inflammatory activation of endothelial cells (20, 21) and phenotypic modulation of SMCs with loss of contractile protein expression (22–24). Inflammatory activation is a likely cause or contributor to SMC phenotypic modulation (25–27). Inflammation and immune function were among the molecular signatures discovered in the *Fbn1^mgR/mgR^* mouse (7). These considerations prompted us to hypothesize that signaling via fibronectin receptors may contribute to the progression of aneurysms.

Our previous work linked pro-inflammatory effects of fibronectin to signaling through the cytoplasmic domain of the integrin α5 subunit(28, 29). These studies developed a chimeric integrin in which the α5 tail was replaced with that of integrin α2, which also pairs with β1 to form integrin α2β1, a receptor for collagen that suppresses inflammatory pathways. This chimeric integrin (denoted α5/2) is fully functional, binds fibronectin and supports cell spreading, cytoskeletal assembly and fibronectin matrix assembly as well as wild-type (WT) α5 but suppresses inflammation similarly to integrin α2(28). Mice in which endogenous α5 is replaced with the chimeric integrin α5/2 are resistant to atherosclerosis, consistent with decreased inflammatory activation within the fibronectin-rich plaque (28, 30). We therefore asked whether the α5/2 mutation would attenuate aortopathy using the mgR model of MFS, in which fibrillin-1 protein expression is decreased by around 85%, resulting in medial pathology with aortic root / ascending aortic aneurysms leading to dissections and rupture that closely resemble human disease (31).

## RESULTS

### Fibronectin accumulates in non-syndromic and Marfan syndrome TAAs

Staining sections of human ascending aorta from healthy control subjects and both syndromic (Marfan) or non-syndromic TAA revealed accumulation of FN as well as diminished levels of the contractile marker smooth muscle α-actin (SMA) in both types of TAAs (Fig. 1A-C). Ascending aortas from the mgR mouse MFS model confirmed a similar increase in FN in the ascending aorta relative to both WT control and α5/2 aortas, which was largely reversed in α5/2 mgR aortas (Fig. 1D, E). Similar to human TAA, SMA (Fig. 1D, F) and smooth muscle protein 22α (Supplemental Fig. S1A, B) were also lower in mgR aortas relative to both WT and chimeric aortas, whereas α5/2 mgR aortas did not significantly differ from WT. Together, these data suggest that increased FN associates with aneurysmal disease, which is captured well by the mgR mouse model, while the α5/2 mutation reduces FN accumulation and improves SMC contractile marker expression in MFS.

**Figure 1.**
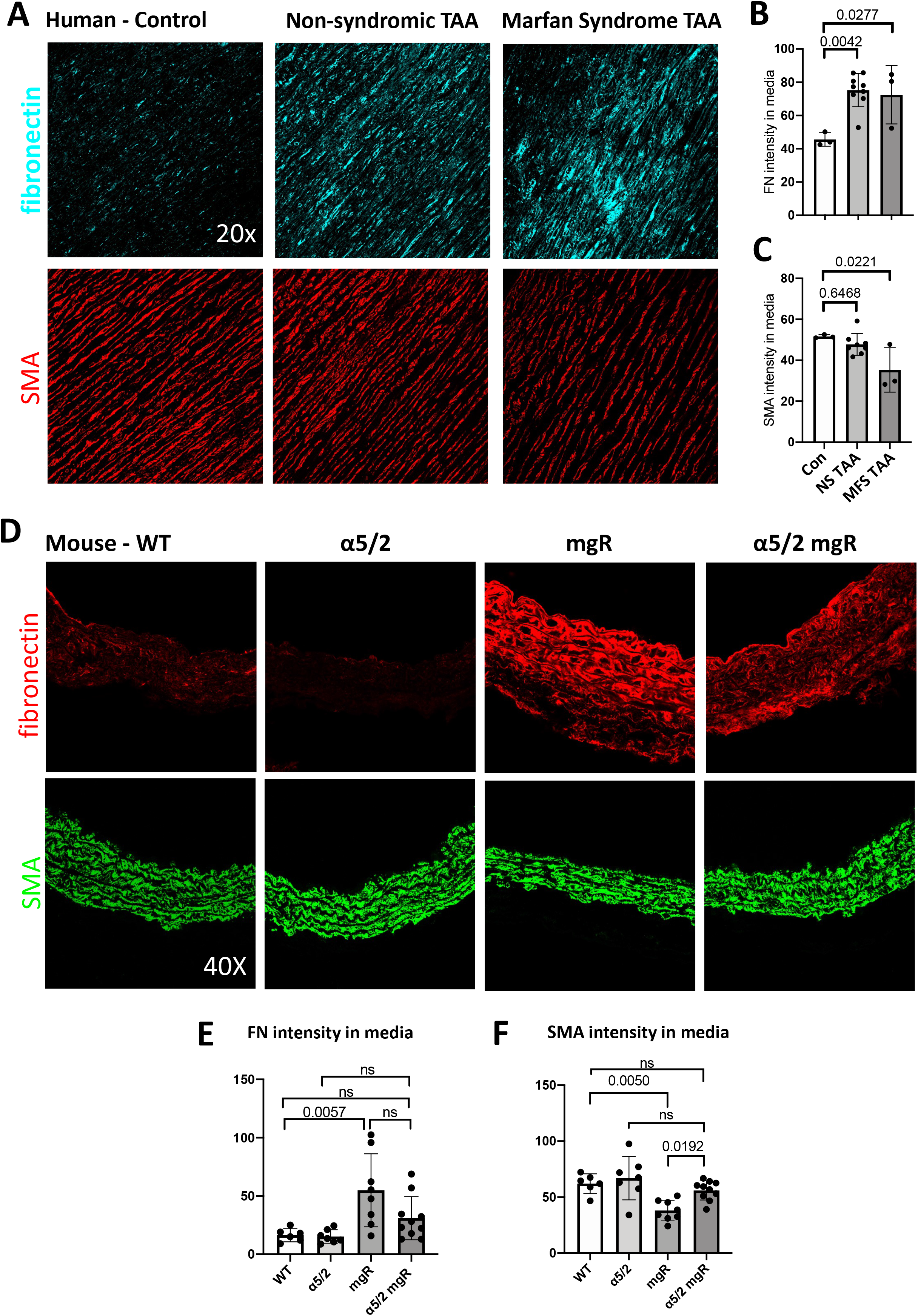
Fibronectin (FN) and smooth muscle α-actin (SMA) in human and murine ascending thoracic aortas. **(A)** Representative human ascending aorta tissue from normal donors, non-syndromic aneurysms and Marfan syndromic patients were stained for FN (top) and SMA (bottom). (**B, C**) Quantification of images in A from human normal controls (Con, n=3), non-syndromic aneurysms (NS TAA, n=9) and Marfan syndrome (MFS TAA, n=3) ascending aortas. (**D**) Ascending aortas from WT mice (n=6), homozygous α5/2 mice (n=7), homozygous mgR mice (n=8), and homozygous α5/2 mgR double mutant mice (n=10) were stained for FN (top) and SMA (bottom). Images were quantified in (**E, F**). Supplemental Fig. S1 shows similar findings for smooth muscle 22α (SM22).

### Integrin α5/2 knock-in limits TAA in mgR mice

We next followed mice from all four groups for up to 90 days after birth. As expected, mgR mice began dying around 40 days of age and lethality reached 60% by 90 days, with death resulting from aortic rupture as reported (31). However, lethality at 90 days was only 15% for the α5/2 mgR mice (Fig. 2A). Given the high mortality of the mgR mice, all subsequent analyses examined 8- to 9-week-old mice. There were no significant differences in body mass (Fig. 2B) or conscious (tail-cuff measured) blood pressure (Fig. 2C) across the four groups at this age. Hence, there was no need for allometric scaling of any dimensioned cardiovascular metrics, and systemic hypertension was not responsible for the increased mortality in the mgR mice.

**Figure 2.**
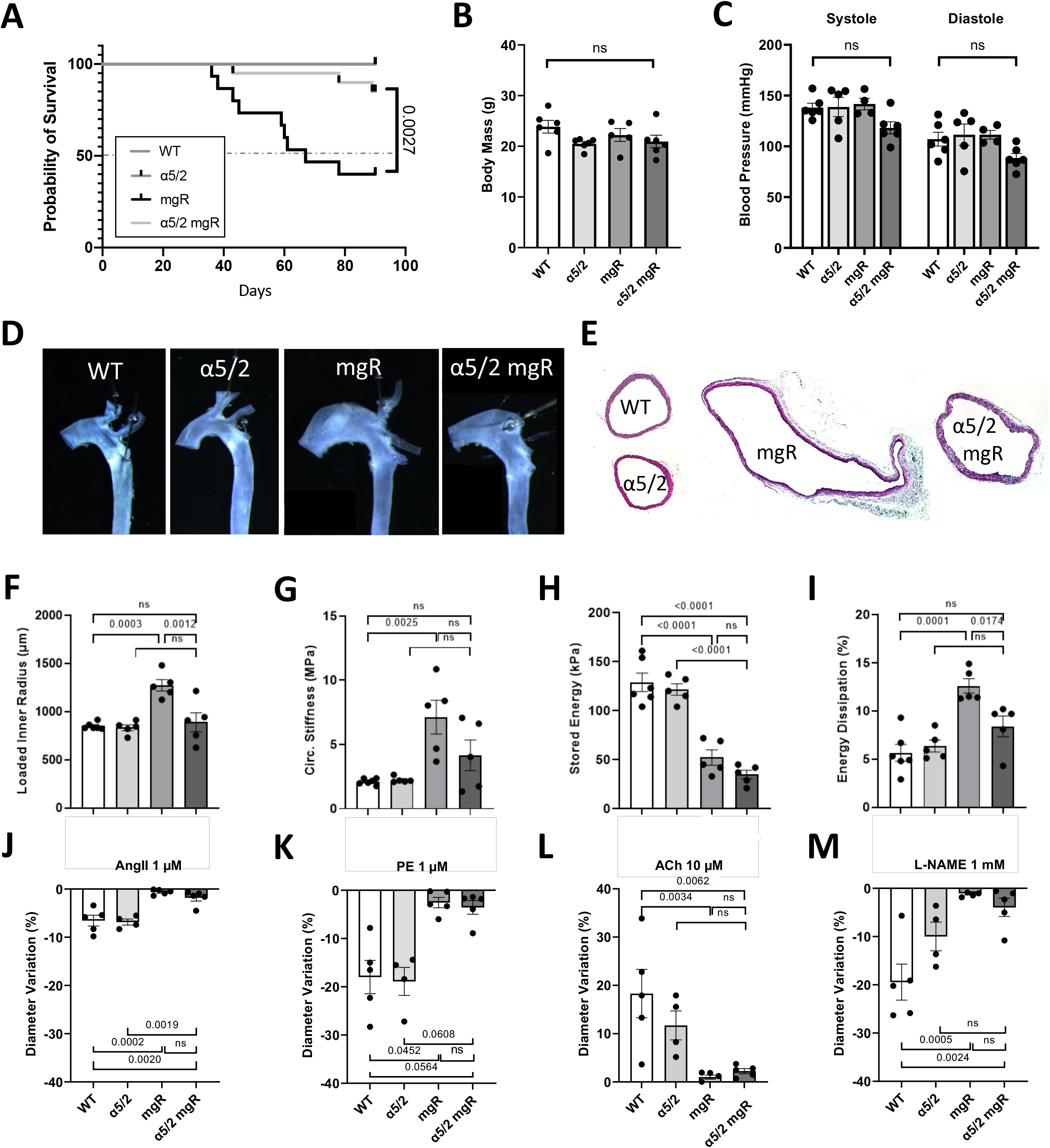
Mouse survival and mechanical parameters in ascending thoracic aortas. WT, α5/2, mgR and mgR α5/2 mice were examined for survival, expansion of the ascending aorta and critical mechanical characteristics: (**A**) Survival over time. (**B**) Body mass and (**C**) mean blood pressure at 8-9 weeks. (**D**) Gross appearance and (**E**) cross section of dissected ascending aortas. (**F-I**) Bulk geometric and passive biomechanical metrics calculated under *ex vivo* equivalent systolic conditions (120 mmHg and specimen-specific *in vivo* axial stretches). (**J-M**) Variation in outer diameter measured during vasoactive biomechanical testing. For panel A: WT n=20, 10 males and 10 females; α5/2, n=20, 10 males and 10 females; mgR n=15, 8 males and 7 females; α5/2 mgR n=20, 10 males and 10 females. For panels B-M, each data point references an individual mouse (WT, n=5-6; α5/2, n=4-6; mgR, n=5; α5/2 mgR, n=5) and columns indicate mean values with standard errors. Statistical bars denote significant differences (paired two-way ANOVA for Fig. 2C, one-way ANOVA for the other panels, with post-hoc Tukey test).

Gross examination *ex vivo* (Fig. 2D) and histological assessment (Figs. 2E, S1C, S2A) of excised aortas showed that the dilatation of the ascending aortas in mgR mice, relative to WT control and α5/2 mice, was dramatically reduced in the α5/2 mgR mice (Fig. S2B). Cross-sectional media area was not significantly affected by the α5/2 substitution in the mgR mice (Fig. S2C), though medial composition was improved significantly (Fig S2D). Consistent with measurements in this unloaded state, luminal diameter of the ascending aorta under simulated *in vivo* systolic loading was significantly greater in the mgR mice (2550 μm, ~1.5 fold increase relative to WT) relative to WT and α5/2 mice (1700 μm); however, lumen diameter was markedly reduced in α5/2 mgR mice (1984 μm, ~1.17-fold increase relative to WT) (Fig. 2F).

Aortic dilatation usually associates with marked changes in bulk mechanical properties, which were quantified *ex vivo* under physiological loading conditions using a custom biaxial testing system). There was little difference in circumferential stretch upon pressurization to 120 mmHg (Fig. S3A) whereas axial stretch was reduced in both mgR aortas and mgR α5/2 ascending aortas with no significant difference between them (Fig. S3B). There was a greater reduction in axial than circumferential wall stress in mgR aortas that was not rescued by the α5/2 substitution (Fig. S3D, E). The most distinguishing mechanical metric in TAAs tends to be increased circumferential material stiffness, which was elevated significantly in the mgR ascending aortas (7.13 MPa) but was not significantly different from WT in the α5/2 mgR mice (Fig. 2G). Axial stiffness was reduced in both mgR and α5/2 mgR relative to WT (Fig. S3C). Although the diminished elastic energy storage capability in mgR was not rescued by the α5/2 substitution (Fig. 2H), the decrease in percent energy dissipation upon cyclic pressurization in the mgR tissue was reversed in mgR α5/2 ascending aortas (Fig. 2I), indicating improved mechanical function. There was, however, no rescue of the vessel-level active vasoconstrictive capacity of the mgR ascending aortas by the α5/2 substitution (Fig. 2J-M, S3F). Full biomechanical analyses for the descending thoracic aorta showed smaller but qualitatively similar results, with functional decline in mgR mice partially rescued by the α5/2 substitution (Fig. S4A-O). Tables S1 and S2 list values of the associated geometric and mechanical metrics for both aortic segments and all four mouse models. Overall, the α5/2 mutation rescues several critical mechanical parameters in the mgR model.

### Elastic laminae damage and collagen remodeling in mgR mice

Compromised elastic fiber integrity is a defining characteristic of the Marfan aorta (5, 31). Multiphoton microscopy under *in vivo* relevant diastolic loading (80 mmHg pressure, specimen-specific *in vivo* values of axial stretch) confirmed marked disruption of the elastic fiber/lamellar structure in the mgR aortas relative to WT controls (overall elastin porosity = 44.3±4.7% (mean±SEM) in mgR vs <3% in both WT and α5/2), with greater porosity in the inner than outer medial layer (Fig. S5A, B); again results were similar, but less marked in the descending thoracic aorta (Fig. S5C). Importantly, this microstructural disruption was markedly reduced in the α5/2 mgR aorta, with an overall elastin porosity = 20.4±2.8% (Fig. 3A, E). Standard unloaded histological cross-sections also showed increased elastin breaks in mgR aortas that was partially rescued in the mgR α5/2 mice (Fig. 3B) in accordance with the changes in elastin amount observed in the unloaded cross-sectioned (Fig. S2A) and loaded wall (Fig. S5A, B). The remaining area fraction in histological cross-sections revealed a significantly higher amount of ground substance (including glycosaminoglycans) in mgR compared to WT mice, with the fraction for α5/2 mgR slightly lower than mgR mice, though still significantly different than WT mice (Fig. S2D). The mgR mice also showed remodeling of adventitial fibrillar collagen, with reduced fiber bundle width which was returned toward normal in mgR α5/2 mice (Fig. 3C). Whereas collagen fiber straightness, volume, and adventitial thickness did not differ across the four groups (Figs 3F and S2E, F), in-plane collagen fiber orientation was significantly more dispersed in mgR compared to WT aortas, with no clear difference among WT, α5/2 and α5/2 mgR aortas (Fig. S2G). Finally, SMC cell density was reduced and adventitial cell density was increased in mgR ascending aorta relative to WT, while the cell densities of α5/2 mgR aorta was closer to WT than mgR aorta (Fig. 3D, G). Microstructure from the descending aorta was evaluated in a similar manner. A reduced severity was observed in the microstructural metrics of the descending aortic region (Fig. S6A-J), wherein mean elastin porosity was 37.2±0.8% in mgR and 16.5±1.2% in α5/2; mgR, with <2.5% in WT and α5/2, and greater in the inner than in the outer layers of the diseased media (Fig. S4C). Elastin porosity thus appears to be an excellent microstructural metric for assigning disease severity in MFS.

**Figure 3.**
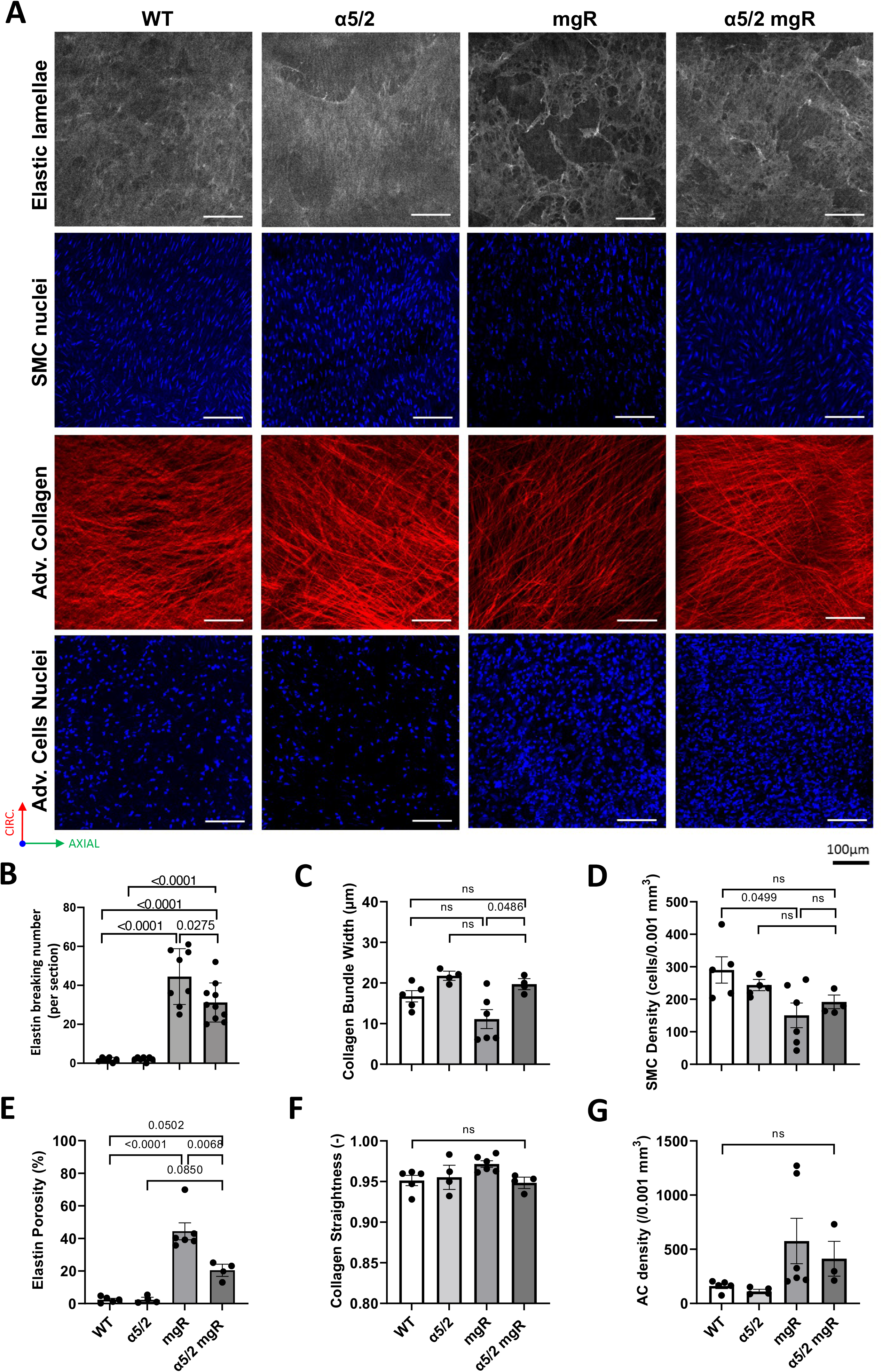
Microstructural characterization of ascending aortas under physiological loading. (**A**) Representative two-photon images acquired at comparable configurations of *ex vivo* equivalent diastolic conditions (80 mmHg and specimen-specific *in vivo* axial stretches) in ascending thoracic aortas from WT, α5/2, mgR and mgR α5/2 mice. Top row: elastin; second row: medial SMC nuclei; third row: adventitial collagen; bottom row: adventitial cell nuclei. (**B**) Number of elastin breaks per histological cross-section fixed at unloaded configuration. (**C-G**) Microstructural metrics quantified from two-photon acquisitions at *ex vivo* equivalent diastolic conditions. Each data point references an individual mouse and columns indicate mean values with standard errors. For Fig. 3B: WT, n=7; α5/2, n=7; mgR, n=8; α5/2 mgR, n=10. For Fig. 3C-G: WT, n=5, α5/2, n=4; mgR, n=6; α5/2 mgR, n=4. Horizontal bars denote statistically significant differences (one-way ANOVA with post-hoc Tukey test).

### Correlations among mechanical metrics and elastin damage

To gain further insights from these multiple mechanical parameters, we calculated linear correlations between key mechanical metrics such as circumferential material stiffness, elastic energy storage, and energy dissipation, increasing luminal diameter and elastic fiber damage of ascending aortas of the four mouse models (Fig. S7). The correlations were particularly strong (p < 0.001) for circumferential stiffness and energy dissipation versus both loaded luminal diameter and volumetric mural damage (elastin porosity), with a clear ordering from no disease (WT and α5/2) to moderate disease (α5/2 mgR) to severe disease (mgR).

### Transcriptomic analyses

We next performed bulk RNAseq of ascending aortic tissue from WT, α5/2, mgR and α5/2 mgR mice at 2 months of age to better understand why the α5/2 substitution improves the mgR phenotype. Principal component analysis separated mgR aortas from all the other 3 groups (Fig. 4D). Consistent with prior studies (32), there were thousands (6544 total, 4024 upregulated, 2520 downregulated) of differentially expressed genes (DEGs) for mgR relative to WT. By contrast, there were fewer DEGs (202) for α5/2 relative to WT and, importantly, the α5/2 substitution in mgR mice largely eliminated differences seen in mgR, with only 315 DEGs for α5/2 mgR relative to WT (Fig. 4A). Functional annotation of DEGs identified changes in gene ontology subcellular processes that are part of inflammatory pathways, which were among the most upregulated in mgR (Fig. 4B). The cytoskeletal/contractile genes, which were among the most downregulated in mgR aorta, were less extensively downregulated in α5/2 mgR (Fig. 4C). Additional pathway enrichment analysis using Gene Ontology Biological Processes for up- and down-regulated genes are show in Figure S8. Note that the aforementioned immunofluorescence results for SMA and SM22 are consistent with the transcriptomic findings (Fig. 1D, F, Fig. S1A, B). The contractile genes *Acta2* (the encoding gene of SMA), *Tagln* (the encoding gene of SM22), as well as *Myh11, Cnn1* and *Itga8,* which is a myocardin regulated protein and thus a surrogate for contractile phenotype, were notably higher in the α5/2 mgR aortas relative to mgR (Fig. 4E), noting that actomyosin function is necessary for both mechanosensing at the cell level and vasoactive control at the tissue level (33). DEGs in mgR relative to WT that were reversed in α5/2 mgR also included cytokines TNF and IL-1β, chemokines CCL3,5 and CXCL13, matrix metalloproteinases MMP9 and MMP25, and their inhibitor TIMP1 in addition to a number of other cytokines, chemokines, and MMPs that were also reduced significantly (Fig. 4F, G). Additional pathway enrichment analysis using Gene Ontology Biological Processes for up- and down-regulated genes.

**Figure 4.**
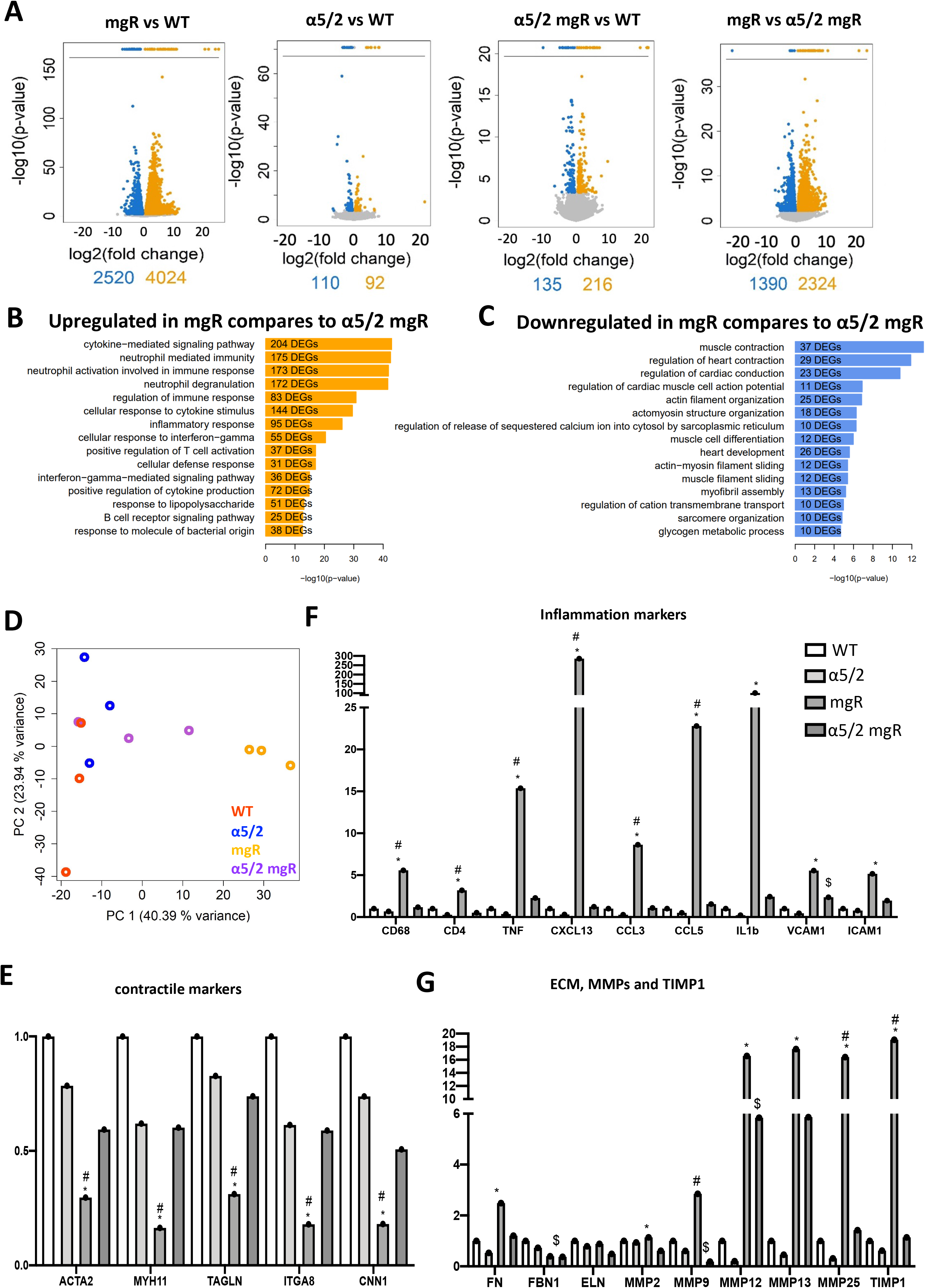
Bulk RNAseq of ascending aortas. (**A**) Volcano plots and pairwise Differentially Expressed Gene (DEG) number comparisons between WT, α5/2, mgR and mgR α5/2 mice, n=3 for each genotype. Significantly down- and up-regulated genes are visualized as blue or orange points, respectively. Numbers below volcano plots indicate total significantly up- and down-regulated genes. (**B-C**) Pathway enrichment analysis using Gene Ontology Biological Processes and Fisher’s Exact Test for up- and down-regulated genes between mgR and α5/2 mgR tissue. (**D**) Log10 gene expression values of all samples were subjected to principal component analysis displayed as Projection of sample gene expression vectors on the first principal component. (**E-G**) Expression of individual inflammatory, contractile and ECM remodeling genes of interest.

### Infiltration of inflammatory cells in non-syndromic and Marfan syndrome TAAs and in vivo

RNAseq results revealed increases in transcripts for CD68 and CD4 in the mgR aortas (Fig. 4F), which were reduced in α5/2 mgR aortas. Immunofluorescence staining confirmed increases in CD68+ (macrophage marker) and CD45+ (pan-leukocyte marker) cells in the ascending aortas in mgR mice relative to control (WT) and α5/2 mice. These increases were substantially reduced in α5/2 mgR aortas (Fig. 5A-D). Whereas CD68+ cells localized mainly in the adventitia in mgR aortas, CD45+ cells distributed across the wall. To test relevance to human aneurysms, we examined sections from non-syndromic TAA and Marfan TAA tissue. CD68 staining in non-syndromic aneurysm tissue showed high variability, consistent with a range of causative factors, but with a trend toward strong elevation, though it did not reach statistical significance (Fig S9A,B). Staining of Marfan tissue showed a weaker trend toward increased CD68-positive cells, again not significant (Fig S9A.C). We also observed a strong trend toward increased CD45-positive cells in non-syndromic aneurysm tissue and a highly significant increase in Marfan tissue relative to healthy controls (Fig. S9D-F).

**Figure 5.**
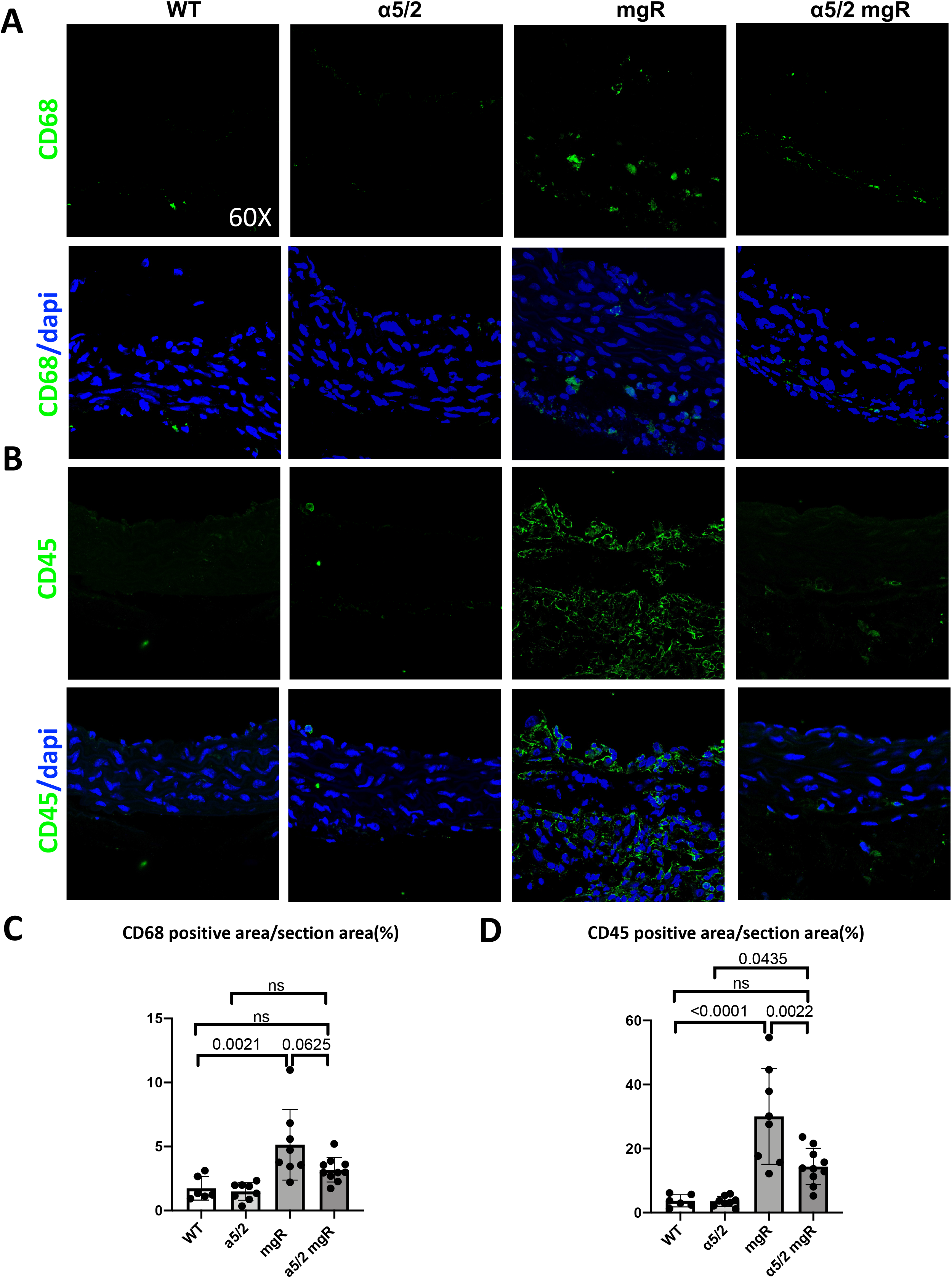
Staining for inflammatory markers in murine ascending thoracic aorta. (**A**) Immunofluorescence of ascending aorta sections from WT, α5/2, mgR and mgR α5/2 mice. Sections were stained for the macrophage marker CD68 or the total hematopoietic marker CD45 together with DAPI to mark cell nuclei as indicated. (**B, C**) Quantification of CD68- and CD45-positive cells per section. Mouse numbers are indicated by individual data points and columns indicate mean values with standard errors. For WT, n=6; α5/2, n=7; mgR, n=8; α5/2 mgR, n=10. Horizontal bars denote statistically significant differences (one-way ANOVA with post-hoc Tukey test).

To further characterize inflammatory activation, we immuno-stained tissue for the p65 subunit of NF-kB and quantified nuclear localization in medial cells as a metric of activation. In human tissue, nuclear p65 was higher in the both Marfan and non-syndromic TAA samples relative to controls (Fig. 6A, B). In mice, nuclear p65 was markedly higher in mgR ascending aortas relative to WT and α5/2 controls but reduced in the α5/2 mgR aortas (Fig. 6C, D). Together, these data confirm increased inflammatory markers in mouse and human TAA, which in the mouse was reversed upon blockade of FN-α5 signaling.

**Figure 6.**
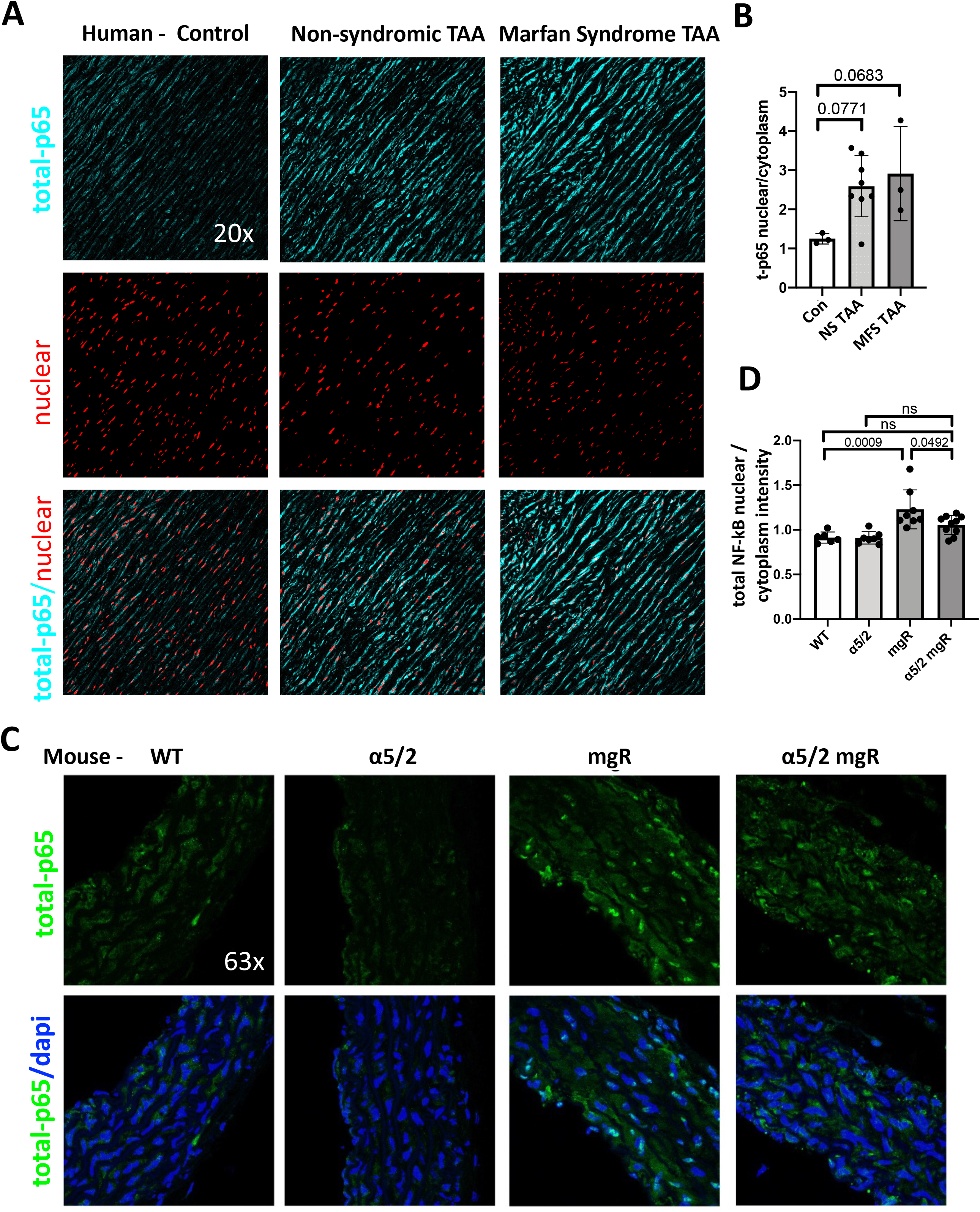
NF-κB in human and murine ascending thoracic aorta. (**A**) Sections from human ascending aortas from healthy controls, non-syndromic aneurysms and Marfan syndromic aneurysms stained for total-p65 plus DAPI as indicated. (**B**) Quantification of nuclear p65 in human ascending aortas: normal donors (Con, n=3), non-syndromic aneurysms (NS TAA, n=9), Marfan syndromic aneurysms (MFS TAA, n=3) (**C**) Ascending aortic sections from WT (n=6), α5/2 (n=7), mgR (n=8) and mgR α5/2 (n=10) mice stained for p65 and DAPI. (**D**) Quantification of nuclear p65 in WT, α5/2, mgR and mgR α5/2 in murine ascending aortic sections. Columns indicate mean values with standard errors. Horizontal bars denote statistically significant differences (one-way ANOVA with post-hoc Tukey test).

### The effect of FN on SMC phenotype in vitro

SMCs play a critical role in the pathogenesis of TAA in Marfan syndrome. To investigate mechanisms, we plated SMCs on FN versus type I collagen *in vitro* to assess direct effects of ECM proteins on SMC phenotype. Aortic SMCs were isolated either from the ascending aorta of a 25kg normal healthy pig or from the entire thoracic aortas of 8- to 9-week-old WT and α5/2 mice. SMCs were plated on 20μg/ml FN or collagen I coated dishes and harvested after 24h. Western blotting showed lower SMA of both porcine (Fig. 7A, B) and WT murine SMCs (Fig. 7D, E) on FN relative to collagen I, which was confirmed by staining for SMA (Fig. 7J, K). By contrast, staining for NF-kB activity by visualization of phospho-p65 revealed a marked increase in SMCs from both species when plated on FN versus collagen I, both by Western blotting (Fig. 7A, C, D, F) and staining (Fig. 7J, L). Importantly, α5/2 murine SMCs neither downregulated SMA nor upregulated phospho-p65 when plated on FN (Fig. 7G-I), confirming that the effects of FN-α5 signaling was abolished in these cells. FN-α5 signaling thus downregulates contractile genes and activates inflammation in SMCs, both are strong contributors to the aneurysmal phenotype.

**Figure 7.**
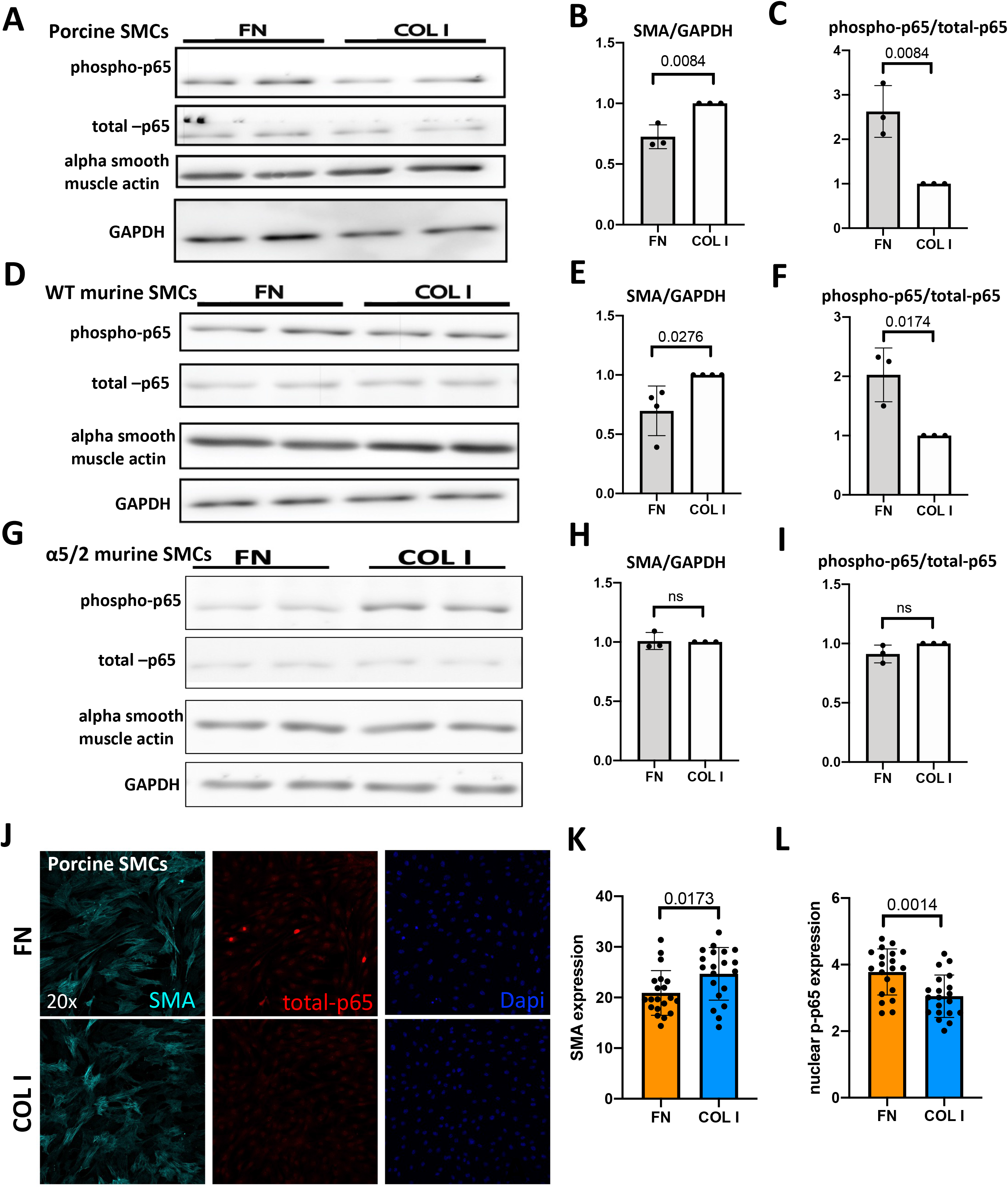
Direct effects of FN on SMCs. (**A**) Porcine SMCs were plated on FN or type I collagen for 24 h then NF-kB activation (p65 phosphorylation) and SMA expression assayed by Western blotting. (**B**) Quantification of SMA results. (**C**) Quantification of p65 results. (**D**) SMCs isolated from WT mice were plated on FN vs type I Coll and NF-kB activation SMA expression assayed as in A. (**E**) Quantification of SMA (**F**) and phospho-p65 (**F**). (**G**) SMCs isolated from α5/2 mice were plated on FN or type I Coll and effects on NF-kB and SMA assayed as in A. Results for SMA quantified in (**H**) and for NF-kB in (**I**). (**J**) Porcine SMC on FN and collagen I were stained for SMA or p65. SMA intensity quantified in (K). Nuclear translocation of p65 quantified in (L). Each dot corresponds to an individual cell. For Fig.7B-I: FN, n=3-4 and COL I, n=3-4; for Fig. 7K-L: n=20. Statistical analysis used twotailed Student’s t-test.

## DISCUSSION

Thoracic aortopathy in MFS is typically considered a disease of the medial layer, but it is increasingly evident that endothelial, smooth muscle, and fibroblasts within the intimal, medial, and adventitial layers, respectively, contribute to pathogenesis (34–36). Hence, rather than focusing on a single cell type, we used a global mouse model that blocked FN-specific inflammatory signaling. Replacing the cytoplasmic domain of the α5 integrin subunit with that of α2 reduces inflammation and the associated atherosclerotic plaque burden in *Apoe^-/-^* mice (28, 30). Elevated FN expression in aneurysms together with recent RNA sequencing and proteomic studies highlighting the importance of integrin signaling and inflammatory processes in TAAs (7, 8, 37) motivated the current study. Our major finding is that the integrin α5/2 mutation substantially protected MFS mice from aortic rupture and death, elastic degradation, and multiple features of mechanical dysfunction in the MFS model. These effects correlated with reduced inflammatory status of the aorta, which is a known driver of ECM degradation and SMC phenotypic switching in aneurysms.

Based on previous work demonstrating the importance of assessing multiple geometric and mechanical metrics (38), we analyzed these parameters in detail. MFS increases structural stiffness, resulting in increased pulse wave velocity and thus central pulse pressure loading of the vulnerable proximal segment (39, 40). More recent phenotyping of aortas from mouse models suggests that the most characteristic biomechanical change in MFS is progressively increased circumferential material stiffness, with values increasing from around 2 MPa in normal murine aortas to 7-11 MPa when aneurysmal, depending on disease severity (41, 42). This increase was partially but significantly blunted in α5/2 mgR aortas, as was the reduced energy dissipation upon cyclic loading (likely due, in part, to the reduced accumulation of ground substance within the wall). Taken together, these findings confirm that signaling through the FN-α5 integrin axis is an important component of the biomechanical phenotype in MFS. Rescue was by no means complete, however, as the α5/2 mutation had little effect on recovering axial stretch and elastically stored energy. The latter is thought to be critical as energy storage during systolic distension enables the aortic wall to recoil elastically during diastole and thus augment blood flow.

There are few detailed biaxial biomechanical data to which to compare these results. Most relevant, a mgR *Ltbp3^-/-^* double mutant has shown the best attenuation of aortopathy to date (7). In that case, the allometrically scaled luminal diameter (noting that *Ltbp3* deletion reduces body mass) was restored toward normal as was circumferential material stiffness (3.06±0.56 MPa), although similar to the present study, axial stretch was not restored and elastically stored energy improved only modestly (43). Reduced elastic energy storage typically results from either compromised elastic fiber integrity or accumulation of other extracellular matrix constituents that prevent competent elastic fibers from deforming, or both. Elastic fiber integrity was improved in both the mgR *Ltbp3^-/-^* and the present mgR α5/2 aorta (Figs. 2, S2, 5). A likely explanation for the low elastic energy storage in α5/2 mgR and mgR *Ltbp3^-/-^* aortas is the lower *in vivo* axial stretch.

Studies using ring myography reported diminished contractile strength in the *Fbn1^C1041G/+^* mouse model (44, 45), as did biaxial testing of the common carotid artery from mgR mice (46). We observed decreased SMC contractile gene and protein expression in the mgR aortas, which was partially corrected in α5/2 mgR. However, this cell-level improvement in contractile markers did not improve vessel-level vasocontractility. We suggest this apparent discrepancy is due in part to an inability to transfer cell-level actomyosin activity through the compromised ECM (47). Related to this, mechano-regulation of ECM requires both actomyosin activity and appropriate connections between the intramural cells and ECM. Circumferential material stiffness, which is critical for artery homeostasis and among the variables that are efficiently restored in α5/2 mgR mice, is highly mechano-regulated (41), suggesting that the improvement in contractile markers in α5/2 mgR may have facilitated this. Inflammation is yet a primary cause of disrupted mechanical homeostasis in disease (47). Taken together, these results suggest that increased inflammatory signaling through FN-α5 adversely affects actomyosin activity in MFS and cellular mechano-sensing, which governs ECM remodeling, consistent with tissue level mechanics not restored.

Consistent with results in human TAAs ((19) and Fig. 1), FN was markedly increased in mgR aortas, but less so in α5/2 mgR mice. The α5/2 chimera is fully competent in FN matrix assembly (28), but FN expression and matrix assembly are regulated by intracellular signaling pathways including those that govern inflammation (48, 49). Our *in vitro* analysis showed that SMC binding to FN decreased smooth muscle cell contractile markers and increased signaling through NF-kB, confirming a pro-inflammatory role for FN in SMCs. Transcriptomic analyses and tissue immunostaining confirmed elevated inflammation in MFS, which was substantially corrected by the α5/2 substitution. Inflammation is a major driver of expression and activation of the MMPs that mediate ECM degradation, including reduced integrity of elastic laminae (50). Reduced inflammation in the α5/2 mice thus likely mediates the improved outcomes. Indeed, deletion of NF-kB/RelA (p65) in mesenchymal cells protects against angiotensin II-induced aortopathy (51), while deletion of MMP2 in the mgR model also reduces elastic fiber degradation and improves lifespans (52). Importantly, NF-kB is among the transcription factors that drive FN gene expression (53, 54), potentially creating a positive feedback loop. Such positive feedback mechanisms are important drivers of progressive illness. Thus, the reduced FN accumulation is likely both a cause and a consequence of lower inflammatory status.

Although the aortic root and ascending aorta are most vulnerable in MFS, the descending thoracic aorta can become increasingly vulnerable in MFS patients following proximal aortic graft replacement (55). This distal vulnerability may result from graft-related alterations in biomechanical loading, from an extended lifespan that allows non-resected segments to deteriorate, or both. Loss of elastic fibers in the descending aorta of mgR mouse is less dramatic than in the ascending but highly significant (56), which is supported by biomechanical assessments (57). The improved biomechanics in α5/2 mgR descending aortas thus supports the general importance of FN-α5 signaling in medial degeneration.

In summary, inflammatory signaling through the FN-integrin α5 axis contributes strongly to the MFS aortic phenotype. Inflammatory activation in the vessel wall due to fibrillin-1 insufficiency results in increased mural FN, which, via α5 – NF-kB signaling, drives increases in chemokines, cytokines, infiltrating immune (CD45+ and CD68+) cells, MMP activity, and further FN deposition. These processes are likely self-amplifying, leading to diminished cell-level mechano-regulation of extracellular matrix and associated compromised wall properties, reduced vessel-level vasoactivity, and reduced wall strength predisposing to aortic rupture. Blocking adverse FN-α5 signaling thus warrants further investigation as a therapeutic target in Marfan syndrome.

## METHODS

### Human samples Immunofluorescence

3 non-diseased aortas from organ donors, 9 non-syndromic TAA and 3 Marfan Syndrome TAA samples were provided by the Tellides lab and Assi lab. Research protocols were approved by the Institutional Review Boards of Yale University and the New England Organ Bank. Segments of ascending aorta was fixed in formalin and embedded in paraffin. Blocks were sectioned at 5um intervals using a paraffin microtome by Yale’s Research Histology Laboratory. Paraffin sections were dewaxed in xylene, boiled for 40 minutes in antigen unmasking solution, citrate-based (H-3300-250, Vector) for antigen retrieval and rehydration. And washed 3 times with Tris-buffered saline (TBS), tissue sections were incubated with primary antibodies diluted in blocking solution (TBS with 3% BSA) overnight at 4 *°C* in a humidified chamber. Antibodies used for human paraffin section staining includes (1:1000, SMA, 50-9760-82, Thermo Fischer, eFluor 660), FN (1:1000, F3648, Sigma), total-p65 (1:500, 4764s, Cell Signaling), CD45 (1:500, 555480, BD science) and CD68 (1:500, ab955, Abcam), dye used for nuclear staining was propidium iodide (1:200, BMS500PI, Thermo Fischer). Sections were washed 3 times with TBS, then incubated with Alexa Fluor 647-conjugated secondary antibody diluted in blocking solution for 1 h at room temperature, washed 3 times in TBS and mounted with Fluoromount-G (0100-01, SouthernBiotech) and imaged with an inverted SP8 confocal microscope (Leica). Image analysis was performed using Fiji software (https://imagej.net/software/fiji/).

### Animals

C57BL/6J mice were purchased from the Jackson laboratory (stock no. 005680), herein as WT mice. *Fbn1^mgR/+^* (mgR) mice (58) were a gift from George Tellides and Francisco Ramirez (available from the Jackson Laboratory as stock no. 005704). The integrin α5/2 chimera C57BL/6 strain was generated using homologous recombination by OZgene (Australia); Floxed-Neo mice were crossed with the CMV-Cre line (stock no. 006054) to create the chimera global knock-in mice as previously described (59), herein as α5/2 mice. α5/2 mice were bred with mgR heterozygous to yield double homozygous mutants, denoted herein as α5/2 mgR mice. All mouse strains used in this paper were on a C57BL/6J background. Mice were maintained in a light- and temperature-controlled environment with free access to food and water. Except survival study, all experiments used 8-9 weeks old male mice. Consistent with federal guidelines, all animal usage was approved by the Yale University Institutional Care and Use Committee.

### Blood pressure

Conscious mice underwent measurement of the systolic and diastolic arterial blood pressures using a tail-cuff device.

### Mouse tissue Immunofluorescence, histology, and morphometric analysis

Mice were euthanized with an overdose of isoflurane (11695-6677-2, Henry Schein) and perfused with PBS through the left ventricle. The ascending aorta (from the aortic root to the brachiocephalic branch) was dissected, and the loose perivascular tissue was gently removed. The mouse ascending aortas were fixed in 3.7% formaldehyde overnight at 4°C, washed with PBS for 3 times, dehydrated in 30% sucrose in PBS for 1 hour and half 30% sucrose half optimal cutting temperature (OCT) for 1 hour, then embedded in OCT and frozen on Dry ice. Tissue blocks were sectioned at 8um. HE staining was done by Yale’s Research Histology Laboratory using standard techniques. HE images were acquired by a Nikon 80i microscope. Morphometry was performed using Fiji. Lumen perimeter was calculated by outlining the internal elastic laminae. Media area was calculated as the area between internal elastic laminae and outer elastic laminae. Frozen sections were incubated with primary antibodies diluted in blocking solution (PBS with 3% BSA), primary antibodies used here included alpha smooth muscle actin (1:1000, SMA, catalog # 50-9760-82, Thermo Fischer, eFluor 660), SM22 (1:500, ab14106, Abcam), FN (1:1000, F3648, Sigma), CD45 (1:500, Af114, R&D), CD68 (1:500, sc6474, Santa Cruz) and total-p65 (1:500, 4764s, Cell Signaling), and a dye: Alexa fluor 633 Hydrazide (A30634, Invitrogen). Detection of unconjugated primary antibodies was visualized with Alexa Fluor 568-, or 647 conjugated IgG (Invitrogen). Sections were mounted with DAPI Fluoromount-G (with Dapi, 0100-20, SouthernbBiotech) and imaged with an inverted SP8 confocal microscope (Leica). Elastin breaking number was calculated from hydrazide-stained sections. Image analysis was performed using Fiji software.

### Biomechanical Phenotyping

Mice were euthanized with an intraperitoneal injection of pentobarbital sodium and phenytoin sodium (Beuthanasia-D; 150 mg/kg). Both the ascending aorta (from the aortic root to the brachiocephalic branch) and the descending thoracic aorta (from the left subclavian to the third intercostal branch) were harvested and loose perivascular tissue was gently removed. Standardized protocols (60, 61) were used to biomechanically phenotype the ascending and descending thoracic aorta from all four groups of mice: WT, α5/2, mgR, and α5/2 mgR. Each aortic segment was mounted within a custom biaxial testing system in a heated (37°C) and oxygenated (95%/5% O_2_/CO_2_) Krebs-Ringer bicarbonate buffered solution containing 2.5 mM CaCl_2_. After stretching the vessel to near its *in vivo* length and pressurizing it to 90 mmHg, smooth muscle cell and endothelial cell function were assessed by sequential exposures: 100 mM KCl (which depolarizes cell membranes, thus inducing contraction), washout, 1 μM phenylephrine (an α1-adrenergic receptor agonist), then 10 μM acetylcholine (which stimulates endothelial production of nitric oxide), and finally 1 mM L-nitroarginine methyl ester (L-NAME, which blocks endothelial derived nitric oxide synthase). Changes in vessel diameter were measured on-line, as were changes in axial force given the isometric-isobaric constraints.

Next, the fluid bath was changed to calcium free Krebs-Ringer bicarbonate buffered solution and the vessel was subjected to a series of cyclic passive pressure-diameter and axial force-length tests. Specifically, following preconditioning, the vessels were cyclically pressurized from 10-140 mmHg while holding the axial length fixed at the *in vivo* value and, subsequently ±5% of this value, then the vessel was cyclically stretched to the maximum value of force measured in the pressure-diameter test while holding the luminal pressure fixed at 10, 60, 90, and 140 mmHg. Luminal pressure, axial force, outer diameter, and axial length were recorded continuously. Figure S5 shows illustrative descending thoracic aortas from each of the four groups during biaxial testing. An independently validated four-fiber family constitutive model was then used to fit simultaneously the unloading portion of the pressure-diameter-axial force-length data from these seven protocols, thus allowing us to compute and compare sample-specific mechanical metrics, including mean biaxial wall stress, biaxial material stiffness, and elastically stored energy (Tables S1, S2). Another parameter is the percentage of elastic energy that is dissipated during cyclic loading, calculated as the energy difference between the loading and unloading portions normalized by the energy associated with the loading (60).

### Microstructural Characterization

Ascending and descending aortas were then removed from the biaxial testing device, re-cannulated, and placed within a chamber that allowed control of axial stretch and luminal pressure of the sample, immersed in calcium free Krebs-Ringer bicarbonate buffered solution, while acquiring microstructural images. The acquisitions were performed with a two-photon microscope (LaVision BioTec TriMScope) powered by a Titanium-Sapphire laser tuned at 840 nm and equipped with a 20X water immersion objective lens (N.A. 0.95). Second harmonic generation (390-425 nm) images revealed fibrillar collagens while two-photon fluorescence (500-550 nm) images revealed elastin, with cell nuclei captured as well (above 550 nm). Three-dimensional (3D) images were acquired with an in-plane field of view of 500 μm x 500 μm at a consistent anatomical location, where in-plane indicates the axial-circumferential plane of the artery. This location coincides with a central volume of the external curvature in the ascending aorta region and a central volume of the ventral region between the second and third pairs of intercostal arteries in the descending segment. Numerical imaging resolution was 0.48 μm/pixel, while the out-of-plane (radial axis) step size was 1 μm/pixel. Each image was then processed as previously described (62, 63). Key metrics measured were the mean absolute volumes of collagen, elastin, and cell nuclei constituents and the mean thickness of adventitial and medial layers. Metrics quantifying the inplane microstructural organization of adventitial collagen fibers include collagen fiber straightness, collagen fiber bundle width, and primary orientation. Metrics related to cell nuclei are mean density of adventitial and smooth muscle cells by volume of adventitia and media, respectively. The characteristic porous appearance of elastic fibers in MFS tissue was characterized by the percentage of elastin porosity that quantifies the empty volume within the elastic lamellae through the thickness of the media(64).

### RNA sequencing

Mice were euthanized with an overdose of isoflurane (11695-6677-2, Henry Schein) and perfused with PBS through the left ventricle. Mouse ascending aorta was dissected and the loose perivascular tissue was gently removed. The tissue was frozen in liquid nitrogen and then crushed on dry ice. Crushed tissue was immersed in RLT lysis buffer (from RNeasy Mini Kit and DNase Digestion set 217004, Qiagen) and vigorously vortexed. Total RNA was isolated using a RNeasy Mini Kit and DNase Digestion set (217004, Qiagen) according to the manufacture’s protocol. Next generation whole-transcriptome sequencing was performed by Yale Center for Genome Analysis. Reads were aligned to the mouse reference genome GRCm38 using STAR 2.5.4b (PMID: 23104886), followed by mapping of read counts to genomic gene locations using feature counts(65). All genes with zero standard deviation over all samples were removed before the following analysis. We calculated the pairwise Pearson correlation between all sample or gene vectors based on the obtained read counts per gene and sample, followed by hierarchical clustering of sample or gene distances (1 – correlation coefficient)/2) using the R functionality hclust (mehod= “average”). Gene expression matrix was rearranged based on the clustering results. Before visualization we transformed gene expression values into log10(gene expression values). Log10 gene expression matrix (zeros were replace by 0.1 before taking logarithm) was centered and subjected to principal component analysis using the R-functionality ‘prcomp’. Differentially expressed genes were identified using DESeq2 1.32.0(66) using an adjusted p-value cutoff of 0.05 and a minimum log2(fold change) of +/− log2(1.3). Up- and downregulated genes were subjected to pathway enrichment analysis using Gene Ontology Biological Processes(67, 68) (downloaded from the enrichR website(69)) and Fisher’s Exact Test.

### In vitro Cell Culture

Pig ascending aorta was harvested in surgical room, the adventitial was peeled and the endothelial cells were scraped by a blade. The tissue was minced into pieces no greater than 2mm at greatest dimension. 10 of these pieces were explanted to a Petri dish and incubated at 37 *°C* under M199 media (11150-059, Gibco) with 20% fetal calf serum (FBS). SMCs grew out of the explants after 7 days and reached sufficient number for passage in about 14 days. Porcine SMCs were grown in M199 media with 10% FBS and cultured up to passage 5. WT and α5/2 SMCs were obtained from whole mouse thoracic aorta. Mice were euthanized with an overdose of isoflurane (11695-6677-2, Henry Schein) and perfused with PBS through the left ventricle. Thoracic aorta was dissected, put in 200ul collagenase I in HBSS (1mg/ml) for 10 minutes, then the adventitia was removed. The aorta was minced into small pieces, put in collagenase I and elastase in 1ml HBSS (collagenase 1.5 mg/ml, elastase 0.5 mg/ml), incubated at 37 °C for 45 minutes, vortexed for 10 seconds, centrifuged at 1000RMP for 5 minutes. The supernatant was discarded, and the cells were resuspended with DMEM media (11995065, Gibco) with 10% FBS. Mouse SMCs were grown in DMEM media with 10% FBS and cultured up to 2 generations.

To compare the effect of FN and collagen I to SMC, 6-well plates or 8 Well Chamber Glass Slide (80841, Ibidi) were coated with 20ug/ml FN or collagen I (354249, Corning) in PBS at 4°C overnight, washed with PBS for 3 times. Porcine cells were starved overnight in M199 media with 2.5% FBS and mouse cells were starved overnight in DMEM media with 1% FBS before plated on FN/collagen I coated plate/glass. SMCs were cultured on FN/collagen I for 24 hours.

### Western blotting

Porcine or mouse SMC were washed with PBS and extracted in Laemmli sample buffer. Samples were separated by SDS-PAGE and transferred onto nitrocellulose membranes. Membranes were blocked with 5% PBS in TBS-T (1% Tween in TBS) and probed with primary antibodies at 4°C for overnight. The targeting proteins were visualized by HRP-conjugated secondary antibodies and subsequent HRP-luminol reaction.

### Cell Immunofluorescence

SMCs were washed with PBS once, fixed for 10 minutes with 3.7% formaldehyde in PBS. Following fixation, cells were permeabilized with 0.5% Triton X-100 in PBS for 10 minutes were incubated with primary antibody in blocking solution (PBS with 3% BSA) at 4°C overnight, then washed 3 times with PBS and incubated with Alexa Fluor 568- or 647-conjugated secondary antibody diluted in blocking solution for 1 h at room temperature, washed 3 times in PBS and mounted with DAPI Fluoromount-G (0100-20, SouthernBiotech) and imaged with an inverted SP8 confocal microscope (Leica). Image analysis was performed using Fiji software.

### Statistics

All experimental data are displayed as mean±SEM. One-way or two-way analysis of variance (ANOVA) were used, as appropriate, to determine roles of genotype amongst survival, mean geometric, mechanical, and microstructural quantifications for more than two groups. Post-hoc pairwise comparisons were performed using the Tukey’s correction. Unpaired two-tailed Student’s t-test was used when only two groups were statistically compared. Analysis of correlation between sample-specific geometric, mechanical, and microstructural metrics was carried out computing the Pearson correlation coefficient. The level of significance was set as p< 0.05 for all analysis. When lower than or very close to the significance level, p values are reported in the figures, together with linear regression curves with the 95% confidence bands in case of significant correlations. Graphs were created using Graphpad Prism 9.3.

## ACKNOWLEGMENTS

This work was supported by a grant from the US National Institutes of Health (P01 HL134605 to MAS, JDH, GT, and RI). We thank Dr. Roland Assi and his laboratory for invaluable contribution.

## DISCLOSURES

JDH is a member of the Scientific Advisory Board of and paid consultant for Heartflow, Inc., an AI-based company focused on non-invasive assessment of coronary artery disease. The other authors declare no conflicts, financial or otherwise.

## Author contributions

MC, CC, JDH and MAS designed the study. MC, CC, KT, PR and EJ conducted experiments and acquired data. MC and CC analyzed and interpreted data. JDH, GT, RI, FR and MAS supervised the work. JH conducted bulk RNAseq analysis. AH collected human samples. KT drew the schematic. MC, CC, JDH and MAS wrote the manuscript with input from all authors.

## SUPPLEMENTAL FIGURE LEGENDS

**Fig. S1**. **Additional histological characterization ascending aortic wall**.

(**A**) Staining ascending aorta tissue from WT, α5/2, mgR and α5/2 mgR mice for smooth muscle 22α (SM22) as in Fig. 1. (**B**) Quantification of images in (A). (**C**) Sections from ascending aorta cross from the indicated mouse strains were stained with Alexa fluor 633 Hydrazide to visualize elastin. Quantifications of the number of elastin breaks per section are reported in Fig. 3. Each data point references an individual mouse (WT, n=6; α5/2, n=7; mgR, n=8; α5/2 mgR n=10) and columns indicate mean values with standard errors. Horizontal bars denote statistically significant differences (one-way ANOVA with post-hoc Tukey test).

**Fig. S2. Additional histological characterization of the ascending aortic wall**.

(**A**) Bright-field Movat’s Pentachrome staining of ascending aorta from WT, mgR and α5/2 mgR mice and pixel- and color-based analysis to classify constituent area fractions. (**B**) Lumen perimeter and (**C**) medial area in ascending aorta cross sections. (**D**) Calculated medial area fractions for individual constituents from Movat staining analysis: fibrillar collagen φ_c_, elastin φ_e_, cytoplasm φ_m_, ground substance φ_g_ (mainly glycosaminoglycans) and fibrin φ_f_, subject to the constraint that φ_c_ + φ_e_ + φ_m_ + φ_g_ +φ_f_ =1. Each data point references an individual mouse (Fig. S2B, C: WT, n=6; α5/2, n=7; mgR, n=8; α5/2 mgR, n=10; Fig. S2D: WT, n=7; mgR, n=7; α5/2 mgR, n=19) and columns indicate mean values with standard errors. Horizontal bars denote statistically significant differences (paired two-way ANOVA for Fig. S2D and one-way ANOVA for the other panels, with post-hoc Tukey test). Additionally, (**E**) quantified results from main Figure 3B-D were combined to assess vessel wall composition. (**F**) Medial and adventitial thicknesses. (**G**) Orientation of fibrillar collagens in the adventitia and SMC nuclei in the media. The center and radius of the spheres indicate the means and variance of the orientation distributions, respectively. Each data point references an individual mouse (WT, n=5-6; α5/2, n=5; mgR, n=5; α5/2 mgR n=5) and columns indicate mean values with standard errors. Horizontal bars denote statistically significant differences (paired two-way ANOVA for Fig. S2E-F and one-way ANOVA for S2G, with post-hoc Tukey test).

**Fig. S3. Additional mechanical and microstructural characterization of the ascending aortic wall**.

(**A-E**) Circumferential stretch, axial stretch, axial stiffness, biaxial wall stress and axial wall stress calculated under *ex vivo* equivalent systolic conditions (120 mmHg and specimen-specific axial stretches) for WT, α5/2, mgR and mgR α5/2 mouse ascending aortas. (**F**) Vasoconstriction of ascending aortas in response to KCl for all four mouse models.

**Fig. S4. Mechanical analysis of the descending aorta**.

**(A-J)** Bulk geometric and passive biomechanical metrics calculated under *ex vivo* equivalent systolic conditions (120 mmHg and specimen-specific *in vivo* axial stretches) and (**K-O**) active biomechanical metrics for the murine descending thoracic aorta (DTA) from WT, α5/2, mgR and α5/2 mgR mice. Each data point references an individual mouse (WT n=5-6, α5/2 n=5-6, mgR n=5, α5/2 mgR n=5) and columns indicate mean values with standard errors. Horizontal bars denote statistically significant differences (oneway ANOVA with post-hoc Tukey test).

**Fig. S5. Transmural variations in elastic lamellae**.

**(A)** Representative two-photon fluorescence images of elastin from the inner to outer media in WT, mgR, and α5/2 mgR ascending aortas under *ex vivo* equivalent diastolic conditions (80 mmHg and specimen-specific *in vivo* axial stretches). (**B, C**) Quantification of transmural distributions of elastin porosity for the ascending thoracic aorta (ATA) and descending thoracic aorta (DTA). Each dotted curve references an individual mouse (WT, n=5; α5/2, n=3; mgR n=5-6; α5/2 mgR, n=3). Solid lines and shadow areas represent best-fit line and 95% confidence intervals for each group, respectively. Vertical bars denote statistically significant differences between slopes (one-way ANOVA with post-hoc Tukey test).

**Fig. S6. Structure of the descending aorta**.

**(A)** Representative two-photon images acquired at comparable configurations of *ex vivo* equivalent diastolic conditions (80 mmHg and specimen-specific *in vivo* axial stretches) in descending thoracic aorta (DTA) from WT, α5/2, mgR and mgR α5/2 mice. Top row: elastin; second row: medial SMC nuclei; third row: adventitial collagen; bottom row: adventitial cell nuclei. (B-J) Microstructural metrics quantified from two-photon acquisitions at *ex vivo* equivalent diastolic conditions in DTAs. Each data point references an individual mouse (WT, n=4; α5/2, n=4; mgR, n=4; α5/2 mgR, n=4) and columns indicate mean values with standard errors. Horizontal bars denote statistically significant differences (paired two-way ANOVA for Fig. S6B-C and one-way ANOVA for Fig. S6D-J, with post-hoc Tukey test).

**Fig. S7. Linear regression between geometric and mechanical metrics**.

(A-L) Linear regressions between multiple geometric and mechanical metrics and elastin porosity for the ascending aorta were calculated. Each data point references an individual mouse (WT, n=5; α5/2, n=4; mgR, n=5; α5/2 mgR, n=4-5) and columns indicate mean values with standard errors. The best-fit line and 95% confidence intervals (solid and dotted lines, respectively) shown when the slope was either nonzero or showed a nearly significant trend.

**Fig. S8. Additional gene ontology analyses**.

Additional pathway enrichment analysis using Gene Ontology Biological Processes and Fisher’s Exact Test for up- and down-regulated genes. Upregulated pathways are visualized as orange bars and downregulated pathways are visualized as blue bars. n=3 for all groups.

**Figure S9. Staining for inflammatory markers in human ascending thoracic aorta**.

(**A**) Sections from human ascending aortas from healthy controls, non-syndromic (NS) aneurysms and Marfan syndromic (MFS) aneurysms were stained for CD68 and elastin. (**B**) Quantification of percent CD68-positive area in the media of normal donors (Con, n=3) and non-syndromic aneurysms (NS TAA, n=9). (**C**) Quantification of percent CD68-positive area in human ascending aorta media between normal donors (Con, n=3) and Marfan syndromic aneurysms (MFS TAA, n=3). (**D**) Sections from human ascending aortas from healthy controls, non-syndromic aneurysms and Marfan syndromic aneurysms stained for CD45 and elastin. (**E**) Quantification of percent CD45-positive area in the media between normal donors (Con, n=3) and non-syndromic aneurysms (NS TAA, n=9). (**F**) Quantification of percent CD45-positive area in media of normal donors (Con, n=3) and Marfan syndromic aneurysms (MFS TAA, n=3). Columns indicate mean values with standard errors. (Student’s t test for Figs. B, C, E, F).

## TABLES

**Table S1.** Key geometric and mechanical metrics for the ascending thoracic aorta (ATA). Results (mean ± standard error) are provided at unloaded as well as two *in vivo* relevant conditions: the specimen specific value of *in vivo* axial stretch and either a 120 mmHg or 80 mmHg distending pressure. Multiphoton images were collected at the *in vivo* stretch and 80 mmHg, then quantified.

**Table S2.** Key geometric and mechanical metrics for the descending thoracic aorta (DTA). Results (mean ± standard error) are provided at unloaded as well as two *in vivo* relevant conditions: the specimen specific value of *in vivo* axial stretch and either a 120 mmHg or 80 mmHg distending pressure.

---

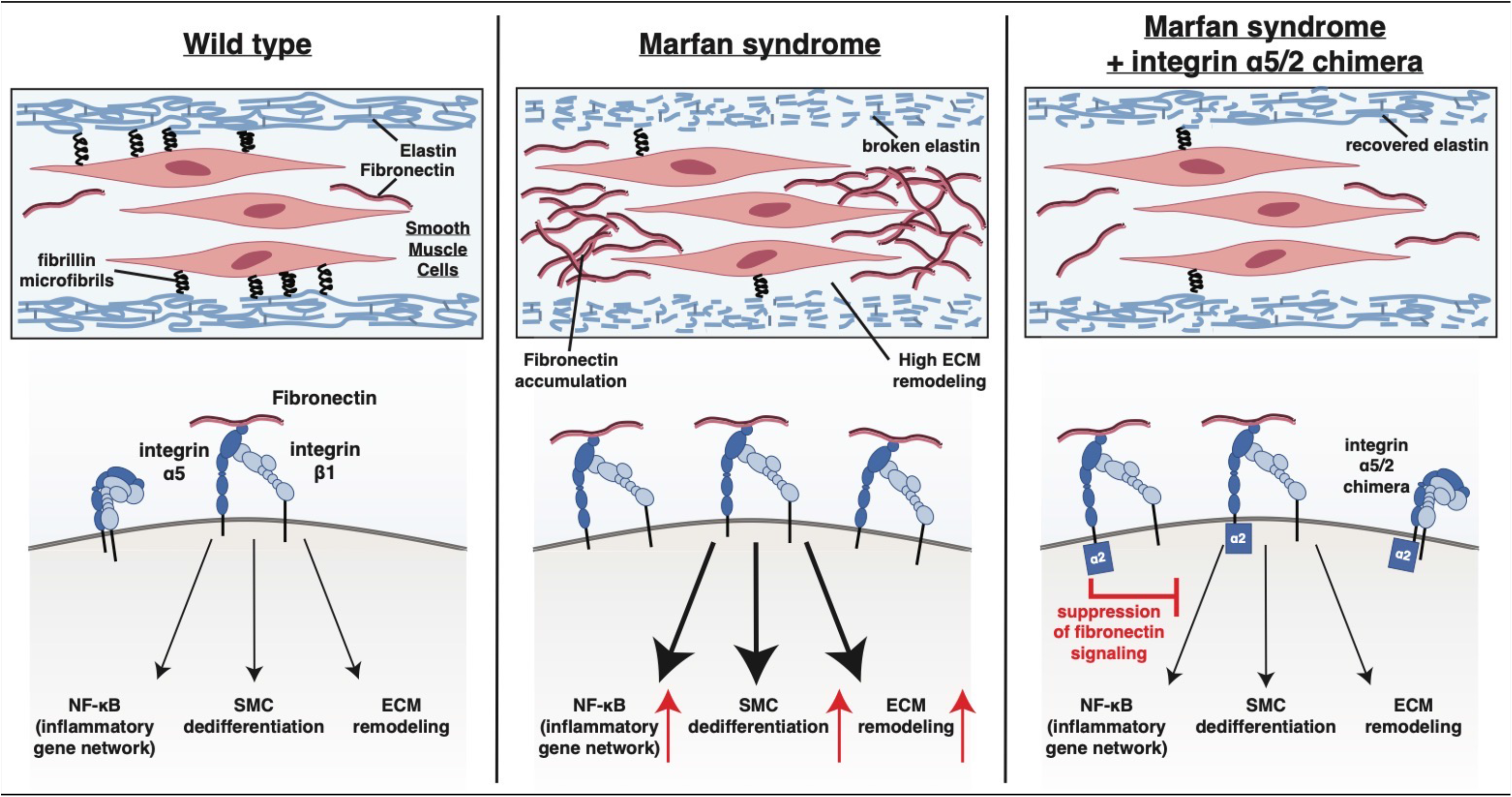
Schematic.

## Notes

### Competing Interest Statement

The authors have declared no competing interest.

### Summary of Updates

Author name

