## supplemental figures and tables for "Fibronectin-integrin α5 signaling promotes thoracic aortic aneurysm in a mouse model of Marfan syndrome"

### Figure S1

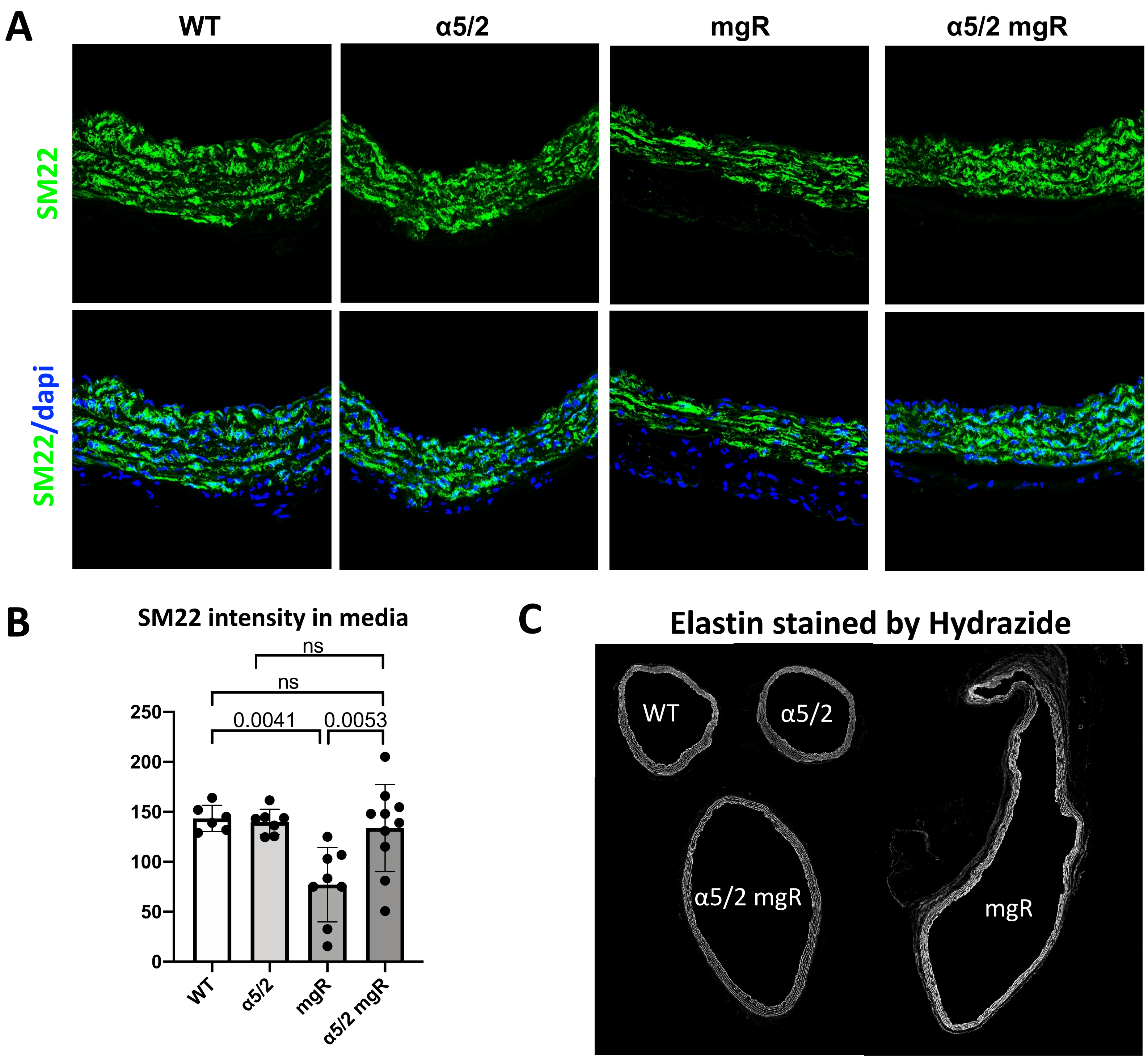

A
WT
α5/2
mgR
α5/2 mgR
SM22
SM22/dapi
B
SM22 intensity in media
ns
ns
0.0041
0.0053
250
200
150
100
50
0
WT
α5/2
mgR
α5/2 mgR
C
Elastin stained by Hydrazide
WT
α5/2
α5/2 mgR
mgR

### Figure S2

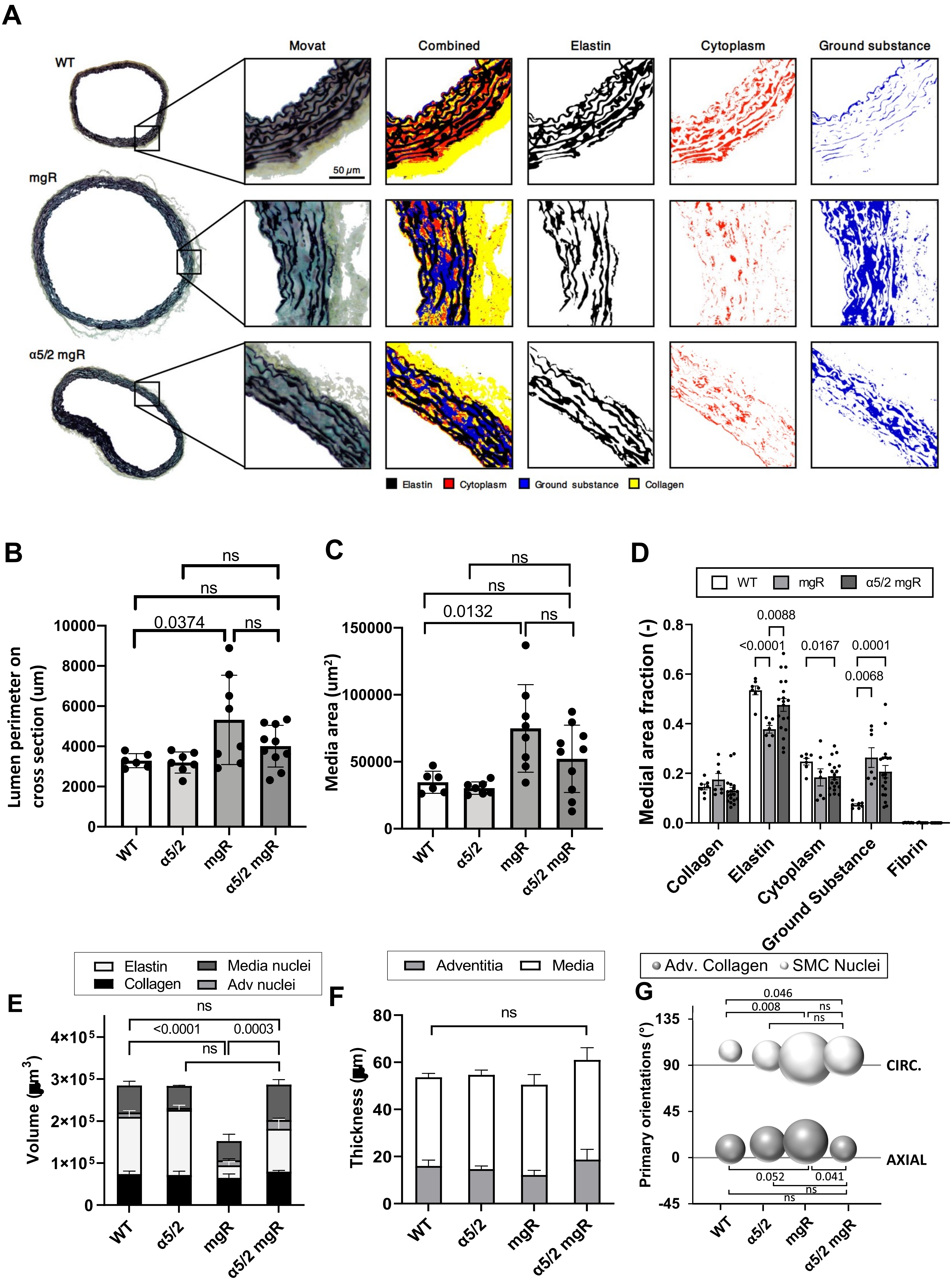

A
WT
mgR
α5/2 mgR
Movat
Combined
Elastin
Cytoplasm
Ground substance
50 μm
Elastin
Cytoplasm
Ground substance
Collagen
B
Lumen perimeter on cross section (um)
0.0374
ns
ns
ns
WT
α5/2
mgR
α5/2 mgR
C
Media area (um²)
0.0132
ns
ns
ns
D
WT
mgR
α5/2 mgR
Medial area fraction (-)
0.0088
<0.0001
0.0167
0.0001
0.0068
Collagen
Elastin
Cytoplasm
Ground Substance
Fibrin
E
Elastin
Collagen
Media nuclei
Adv nuclei
Volume (μm³)
ns
<0.0001
0.0003
ns
F
Adventitia
Media
Thickness (μm)
ns
G
Adv. Collagen
SMC Nuclei
Primary orientations (°)
0.046
0.008
ns
ns
CIRC.
AXIAL
0.052
0.041
ns
ns

### Figure S3

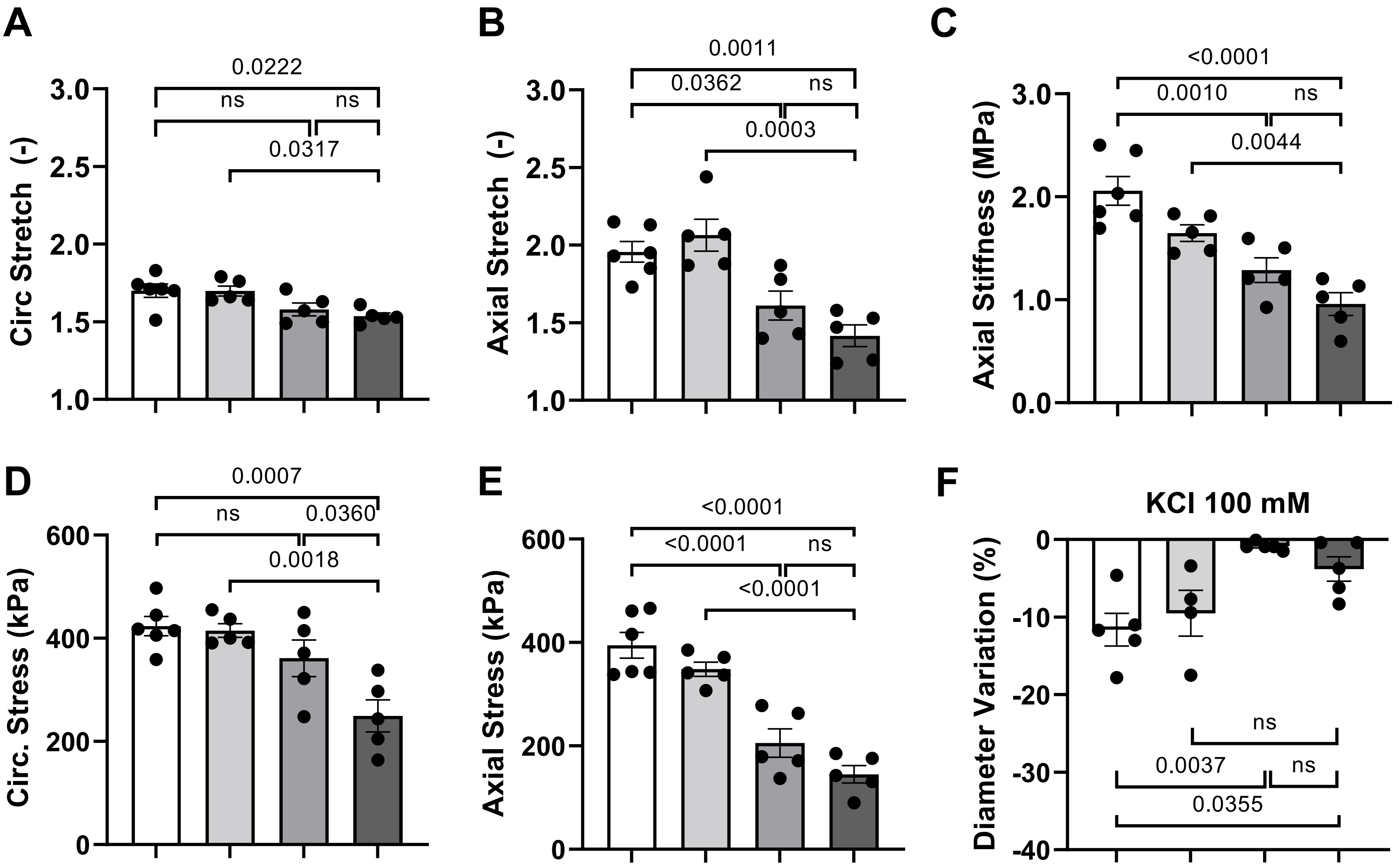

A
Circ Stretch (-)
3.0
2.5
2.0
1.5
1.0
0.0222
ns
ns
0.0317
B
Axial Stretch (-)
0.0011
0.0362
ns
0.0003
C
Axial Stiffness (MPa)
3.0
2.0
1.0
0.0
<0.0001
0.0010
ns
0.0044
D
Circ. Stress (kPa)
600
400
200
0
0.0007
ns
0.0360
0.0018
E
Axial Stress (kPa)
<0.0001
<0.0001
ns
<0.0001
F
KCl 100 mM
Diameter Variation (%)
0
-10
-20
-30
-40
ns
0.0037
ns
0.0355

### Figure S4

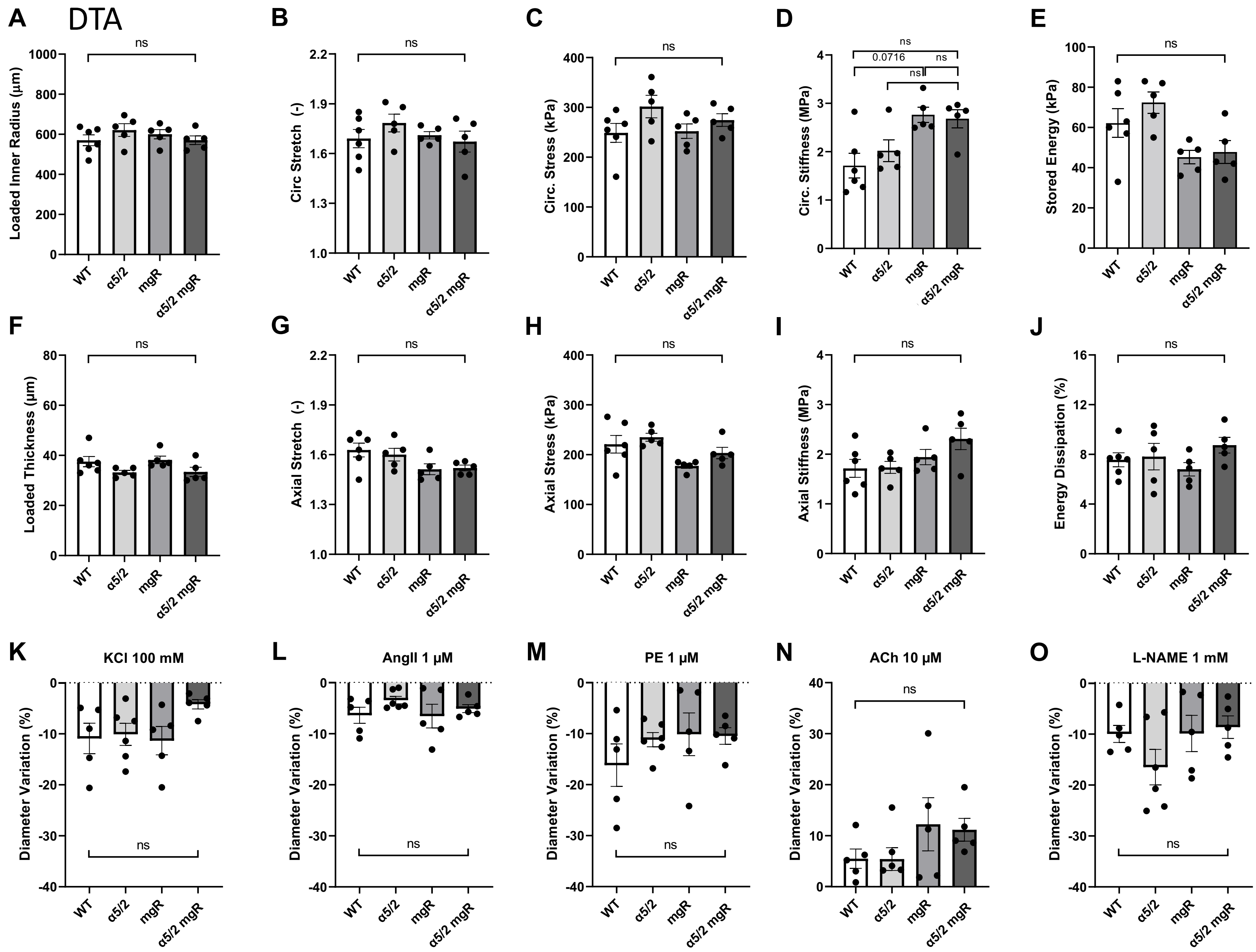
A
DTA
B
C
D
E
F
G
H
I
J
K
L
M
N
O
Loaded Inner Radius (µm)
Circ Stretch (-)
Circ. Stress (kPa)
Circ. Stiffness (MPa)
Stored Energy (kPa)
Loaded Thickness (µm)
Axial Stretch (-)
Axial Stress (kPa)
Axial Stiffness (MPa)
Energy Dissipation (%)
KCl 100 mM
AngII 1 µM
PE 1 µM
ACh 10 µM
L-NAME 1 mM
Diameter Variation (%)
0.0716
ns
WT
α5/2
mgR
α5/2 mgR

### Figure S5

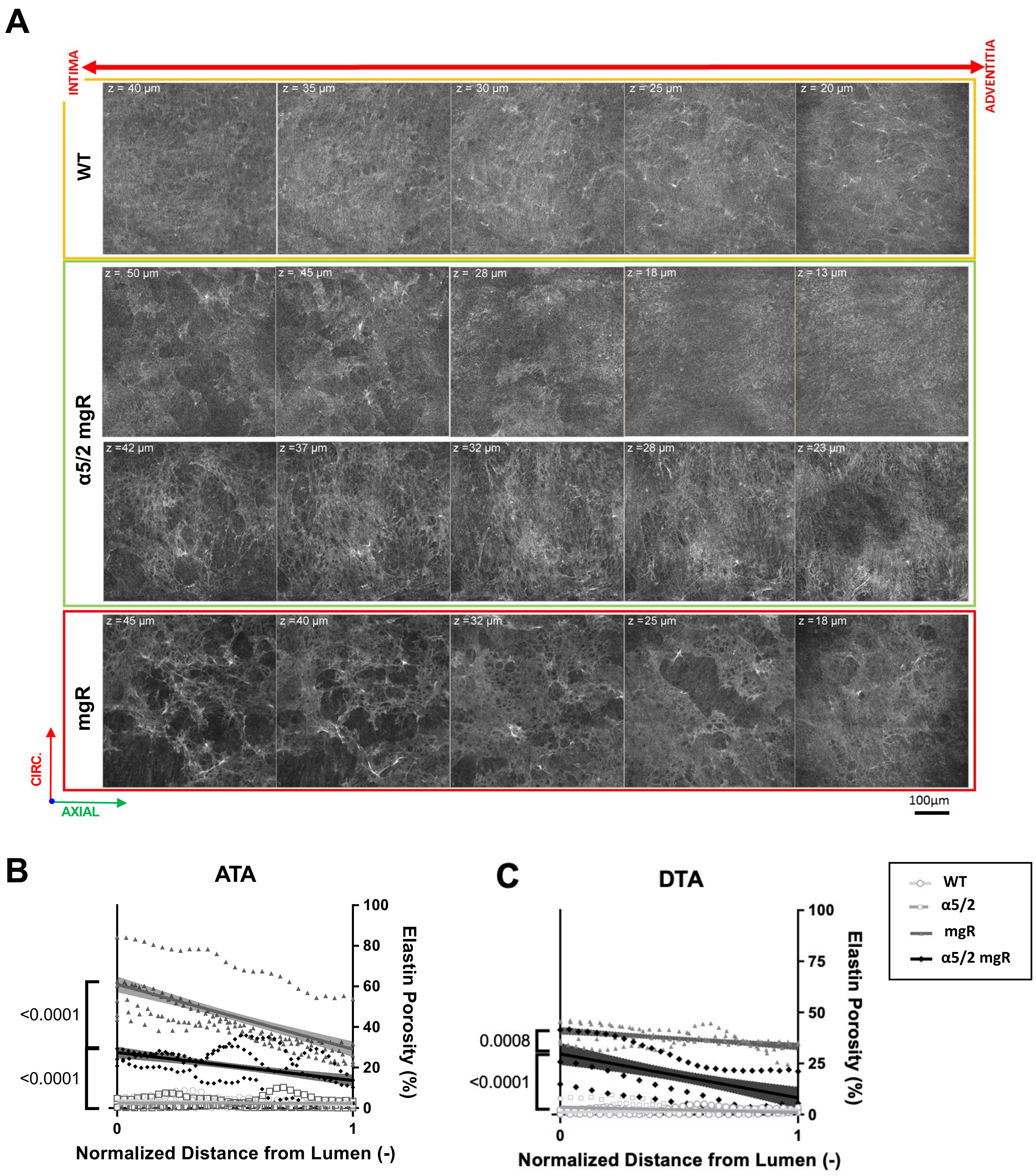

A
INTIMA
ADVENTITIA
WT
z = 40 μm
z = 35 μm
z = 30 μm
z = 25 μm
z = 20 μm
α5/2 mgR
z = 50 μm
z = 45 μm
z = 28 μm
z = 18 μm
z = 13 μm
z =42 μm
z =37 μm
z =32 μm
z =28 μm
z =23 μm
mgR
z =45 μm
z =40 μm
z =32 μm
z =25 μm
z =18 μm
CIRC.
AXIAL
100μm
B
ATA
<0.0001
<0.0001
Elastin Porosity (%)
Normalized Distance from Lumen (-)
C
DTA
0.0008
<0.0001
Elastin Porosity (%)
Normalized Distance from Lumen (-)
WT
α5/2
mgR
α5/2 mgR

### Figure S6

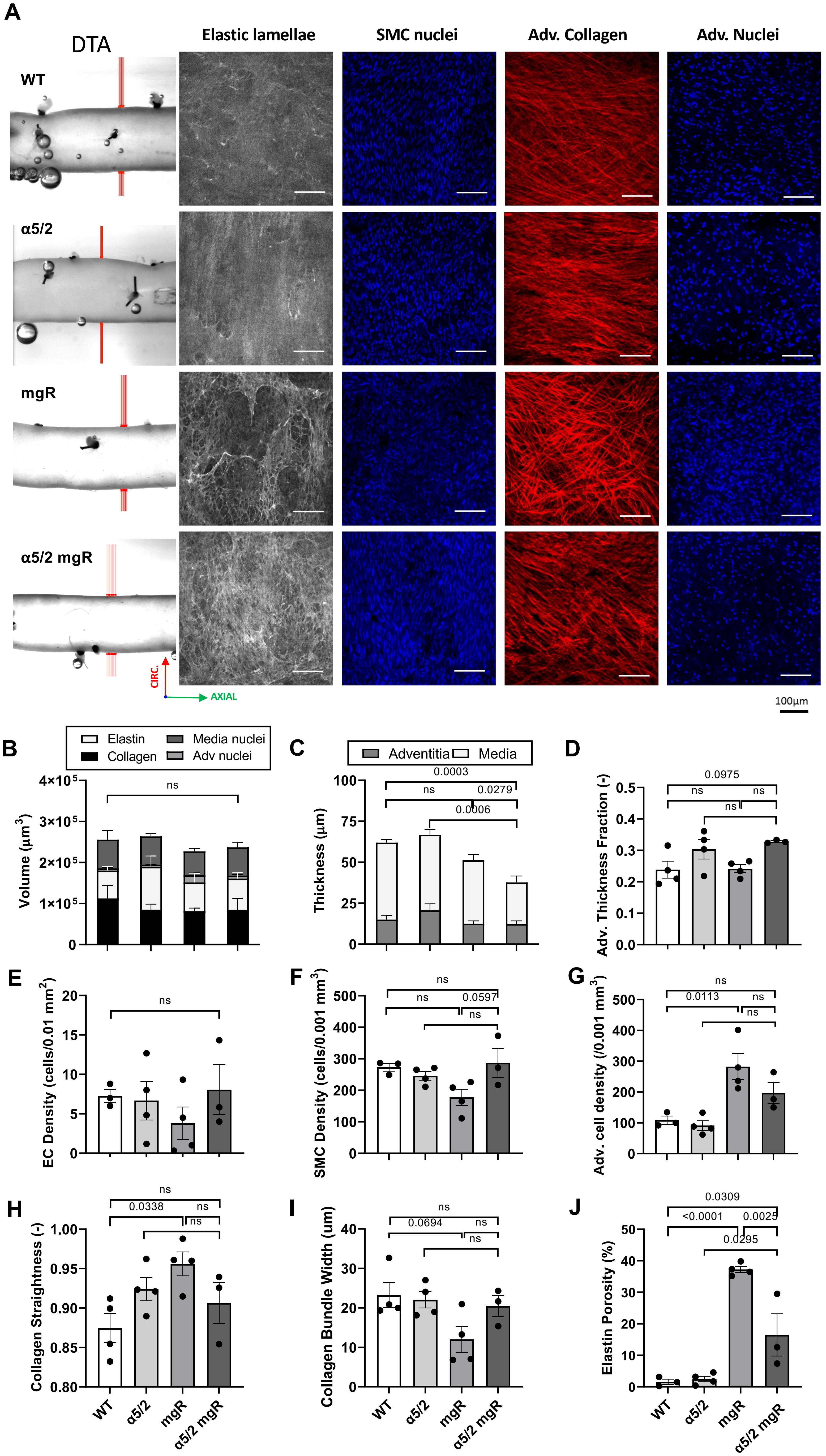

A
DTA
Elastic lamellae
SMC nuclei
Adv. Collagen
Adv. Nuclei
WT
α5/2
mgR
α5/2 mgR
CIRC.
AXIAL
100µm
B
Elastin
Collagen
Media nuclei
Adv nuclei
Volume (µm³)
ns
C
Adventitia
Media
Thickness (µm)
0.0003
ns
0.0279
0.0006
D
Adv. Thickness Fraction (-)
0.0975
ns
ns
ns
E
EC Density (cells/0.01 mm²)
ns
F
SMC Density (cells/0.001 mm³)
ns
ns
0.0597
ns
G
Adv. cell density (/0.001 mm³)
ns
0.0113
ns
ns
H
Collagen Straightness (-)
ns
0.0338
ns
ns
I
Collagen Bundle Width (um)
ns
0.0694
ns
ns
J
Elastin Porosity (%)
0.0309
<0.0001
0.0025
0.0295
WT
α5/2
mgR
α5/2 mgR

### Figure S7

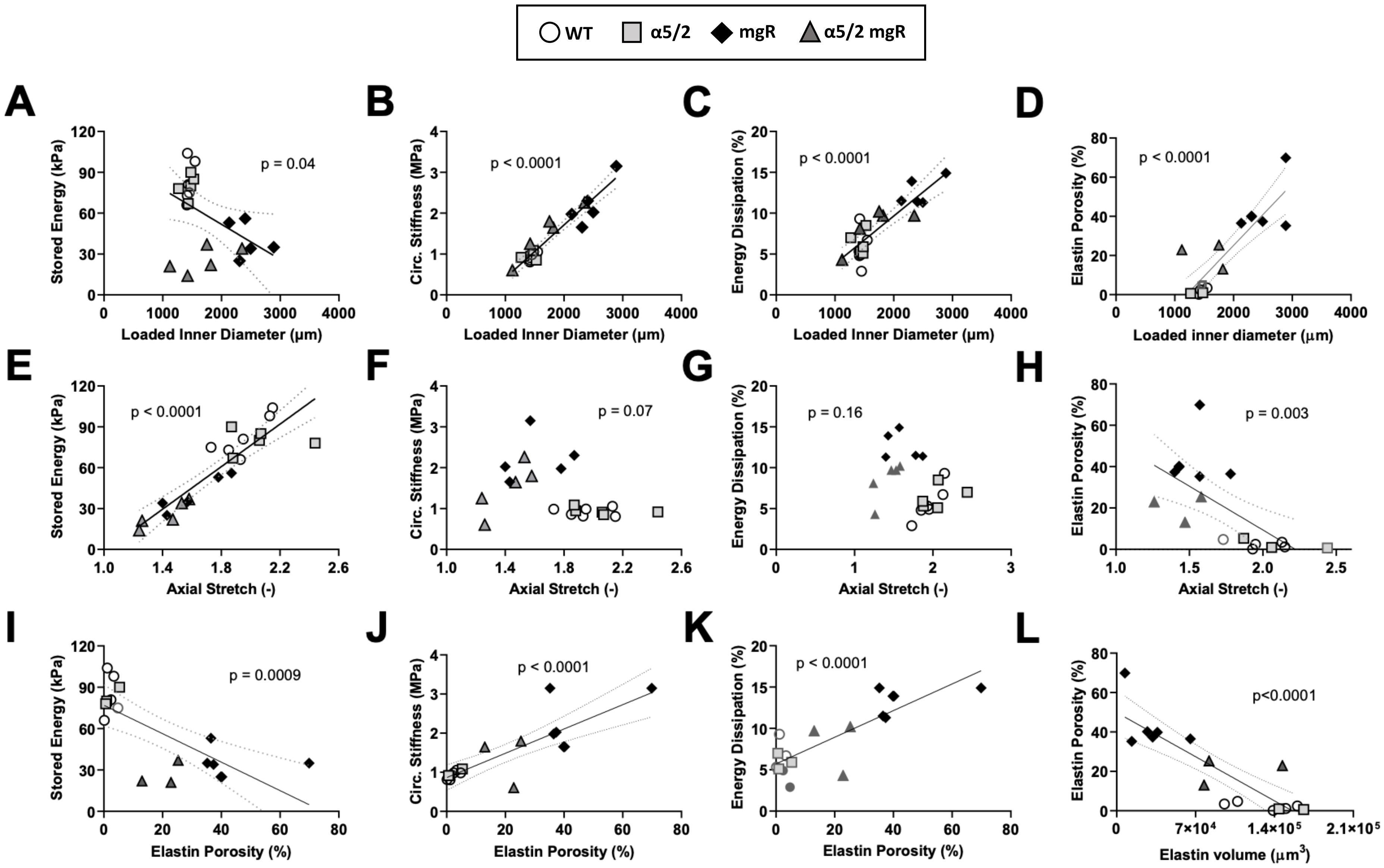
WT
α5/2
mgR
α5/2 mgR
A
Stored Energy (kPa)
Loaded Inner Diameter (µm)
p = 0.04
B
Circ. Stiffness (MPa)
Loaded Inner Diameter (µm)
p < 0.0001
C
Energy Dissipation (%)
Loaded Inner Diameter (µm)
p < 0.0001
D
Elastin Porosity (%)
Loaded inner diameter (µm)
p < 0.0001
E
Stored Energy (kPa)
Axial Stretch (-)
p < 0.0001
F
Circ. Stiffness (MPa)
Axial Stretch (-)
p = 0.07
G
Energy Dissipation (%)
Axial Stretch (-)
p = 0.16
H
Elastin Porosity (%)
Axial Stretch (-)
p = 0.003
I
Stored Energy (kPa)
Elastin Porosity (%)
p = 0.0009
J
Circ. Stiffness (MPa)
Elastin Porosity (%)
p < 0.0001
K
Energy Dissipation (%)
Elastin Porosity (%)
p < 0.0001
L
Elastin Porosity (%)
Elastin volume (µm³)
p<0.0001

### Figure S8

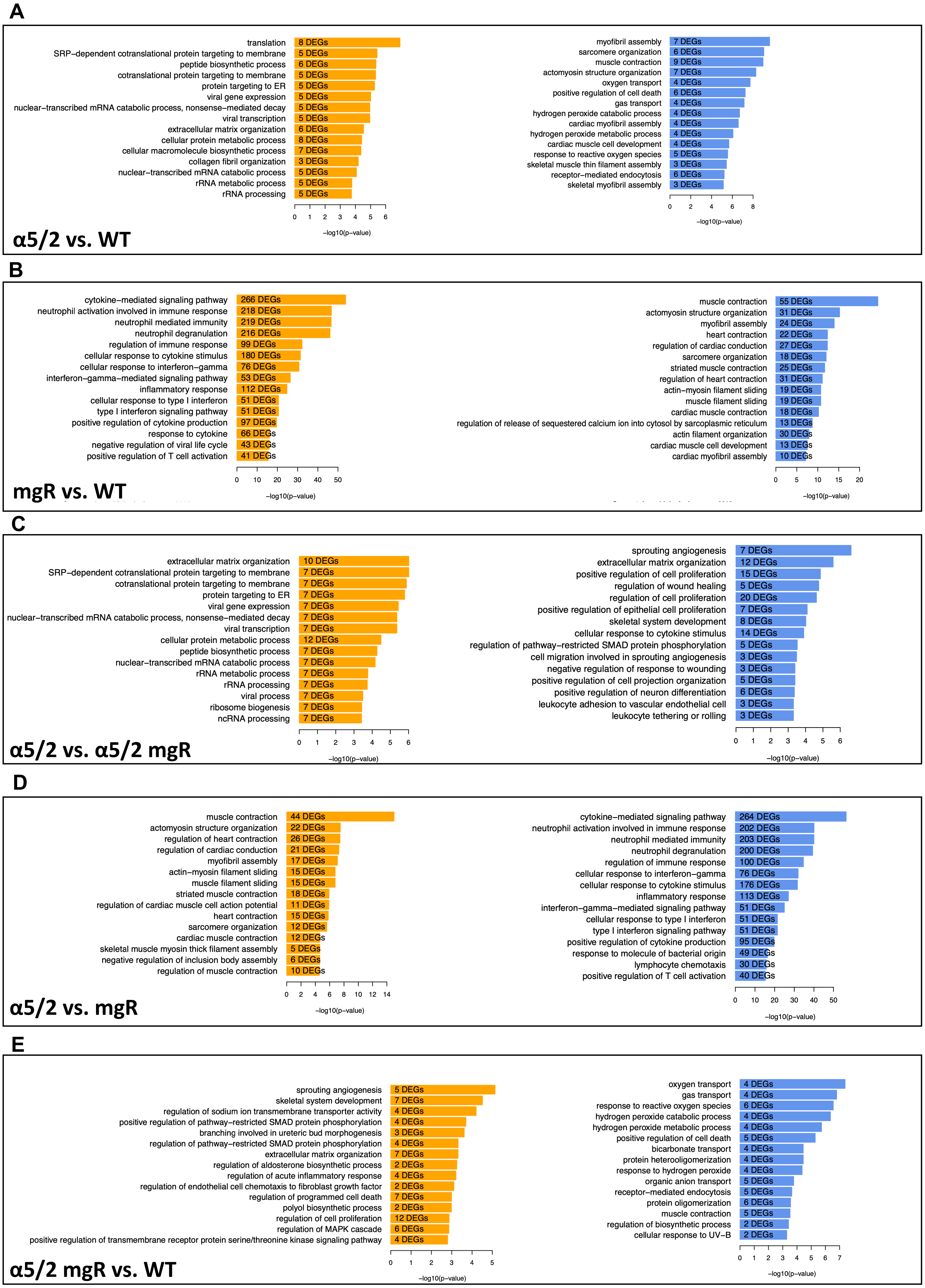
A
translation
SRP−dependent cotranslational protein targeting to membrane
peptide biosynthetic process
cotranslational protein targeting to membrane
protein targeting to ER
viral gene expression
nuclear−transcribed mRNA catabolic process, nonsense−mediated decay
viral transcription
extracellular matrix organization
cellular protein metabolic process
cellular macromolecule biosynthetic process
collagen fibril organization
nuclear−transcribed mRNA catabolic process
rRNA metabolic process
rRNA processing
8 DEGs
5 DEGs
6 DEGs
5 DEGs
5 DEGs
5 DEGs
5 DEGs
5 DEGs
6 DEGs
8 DEGs
7 DEGs
3 DEGs
5 DEGs
5 DEGs
5 DEGs
−log10(p−value)
myofibril assembly
sarcomere organization
muscle contraction
actomyosin structure organization
oxygen transport
positive regulation of cell death
gas transport
hydrogen peroxide catabolic process
cardiac myofibril assembly
hydrogen peroxide metabolic process
cardiac muscle cell development
response to reactive oxygen species
skeletal muscle thin filament assembly
receptor−mediated endocytosis
skeletal myofibril assembly
7 DEGs
6 DEGs
9 DEGs
7 DEGs
4 DEGs
6 DEGs
4 DEGs
4 DEGs
4 DEGs
4 DEGs
4 DEGs
5 DEGs
3 DEGs
6 DEGs
3 DEGs
−log10(p−value)
α5/2 vs. WT
B
cytokine−mediated signaling pathway
neutrophil activation involved in immune response
neutrophil mediated immunity
neutrophil degranulation
regulation of immune response
cellular response to cytokine stimulus
cellular response to interferon−gamma
interferon−gamma−mediated signaling pathway
inflammatory response
cellular response to type I interferon
type I interferon signaling pathway
positive regulation of cytokine production
response to cytokine
negative regulation of viral life cycle
positive regulation of T cell activation
266 DEGs
218 DEGs
219 DEGs
216 DEGs
99 DEGs
180 DEGs
76 DEGs
53 DEGs
112 DEGs
51 DEGs
51 DEGs
97 DEGs
66 DEGs
43 DEGs
41 DEGs
−log10(p−value)
muscle contraction
actomyosin structure organization
myofibril assembly
heart contraction
regulation of cardiac conduction
sarcomere organization
striated muscle contraction
regulation of heart contraction
actin−myosin filament sliding
muscle filament sliding
cardiac muscle contraction
regulation of release of sequestered calcium ion into cytosol by sarcoplasmic reticulum
actin filament organization
cardiac muscle cell development
cardiac myofibril assembly
55 DEGs
31 DEGs
24 DEGs
22 DEGs
27 DEGs
18 DEGs
25 DEGs
31 DEGs
19 DEGs
19 DEGs
18 DEGs
13 DEGs
30 DEGs
13 DEGs
10 DEGs
−log10(p−value)
mgR vs. WT
C
extracellular matrix organization
SRP−dependent cotranslational protein targeting to membrane
cotranslational protein targeting to membrane
protein targeting to ER
viral gene expression
nuclear−transcribed mRNA catabolic process, nonsense−mediated decay
viral transcription
cellular protein metabolic process
peptide biosynthetic process
nuclear−transcribed mRNA catabolic process
rRNA metabolic process
rRNA processing
viral process
ribosome biogenesis
ncRNA processing
10 DEGs
7 DEGs
7 DEGs
7 DEGs
7 DEGs
7 DEGs
7 DEGs
12 DEGs
7 DEGs
7 DEGs
7 DEGs
7 DEGs
7 DEGs
7 DEGs
7 DEGs
−log10(p−value)
sprouting angiogenesis
extracellular matrix organization
positive regulation of cell proliferation
regulation of wound healing
regulation of cell proliferation
positive regulation of epithelial cell proliferation
skeletal system development
cellular response to cytokine stimulus
regulation of pathway−restricted SMAD protein phosphorylation
cell migration involved in sprouting angiogenesis
negative regulation of response to wounding
positive regulation of cell projection organization
positive regulation of neuron differentiation
leukocyte adhesion to vascular endothelial cell
leukocyte tethering or rolling
7 DEGs
12 DEGs
15 DEGs
5 DEGs
20 DEGs
7 DEGs
8 DEGs
14 DEGs
5 DEGs
3 DEGs
3 DEGs
5 DEGs
6 DEGs
3 DEGs
3 DEGs
−log10(p−value)
α5/2 vs. α5/2 mgR
D
muscle contraction
actomyosin structure organization
regulation of heart contraction
regulation of cardiac conduction
myofibril assembly
actin−myosin filament sliding
muscle filament sliding
striated muscle contraction
regulation of cardiac muscle cell action potential
heart contraction
sarcomere organization
cardiac muscle contraction
skeletal muscle myosin thick filament assembly
negative regulation of inclusion body assembly
regulation of muscle contraction
44 DEGs
22 DEGs
26 DEGs
21 DEGs
17 DEGs
15 DEGs
15 DEGs
18 DEGs
11 DEGs
15 DEGs
12 DEGs
12 DEGs
5 DEGs
6 DEGs
10 DEGs
−log10(p−value)
cytokine−mediated signaling pathway
neutrophil activation involved in immune response
neutrophil mediated immunity
neutrophil degranulation
regulation of immune response
cellular response to interferon−gamma
cellular response to cytokine stimulus
inflammatory response
interferon−gamma−mediated signaling pathway
cellular response to type I interferon
type I interferon signaling pathway
positive regulation of cytokine production
response to molecule of bacterial origin
lymphocyte chemotaxis
positive regulation of T cell activation
264 DEGs
202 DEGs
203 DEGs
200 DEGs
100 DEGs
76 DEGs
176 DEGs
113 DEGs
51 DEGs
51 DEGs
51 DEGs
95 DEGs
49 DEGs
30 DEGs
40 DEGs
−log10(p−value)
α5/2 vs. mgR
E
sprouting angiogenesis
skeletal system development
regulation of sodium ion transmembrane transporter activity
positive regulation of pathway−restricted SMAD protein phosphorylation
branching involved in ureteric bud morphogenesis
regulation of pathway−restricted SMAD protein phosphorylation
extracellular matrix organization
regulation of aldosterone biosynthetic process
regulation of acute inflammatory response
regulation of endothelial cell chemotaxis to fibroblast growth factor
regulation of programmed cell death
polyol biosynthetic process
regulation of cell proliferation
regulation of MAPK cascade
positive regulation of transmembrane receptor protein serine/threonine kinase signaling pathway
5 DEGs
7 DEGs
4 DEGs
4 DEGs
3 DEGs
4 DEGs
7 DEGs
2 DEGs
4 DEGs
2 DEGs
7 DEGs
2 DEGs
12 DEGs
6 DEGs
4 DEGs
−log10(p−value)
oxygen transport
gas transport
response to reactive oxygen species
hydrogen peroxide catabolic process
hydrogen peroxide metabolic process
positive regulation of cell death
bicarbonate transport
protein heterooligomerization
response to hydrogen peroxide
organic anion transport
receptor−mediated endocytosis
protein oligomerization
muscle contraction
regulation of biosynthetic process
cellular response to UV−B
4 DEGs
4 DEGs
6 DEGs
4 DEGs
4 DEGs
5 DEGs
4 DEGs
4 DEGs
4 DEGs
5 DEGs
5 DEGs
6 DEGs
5 DEGs
2 DEGs
2 DEGs
−log10(p−value)
α5/2 mgR vs. WT

### Figure S9

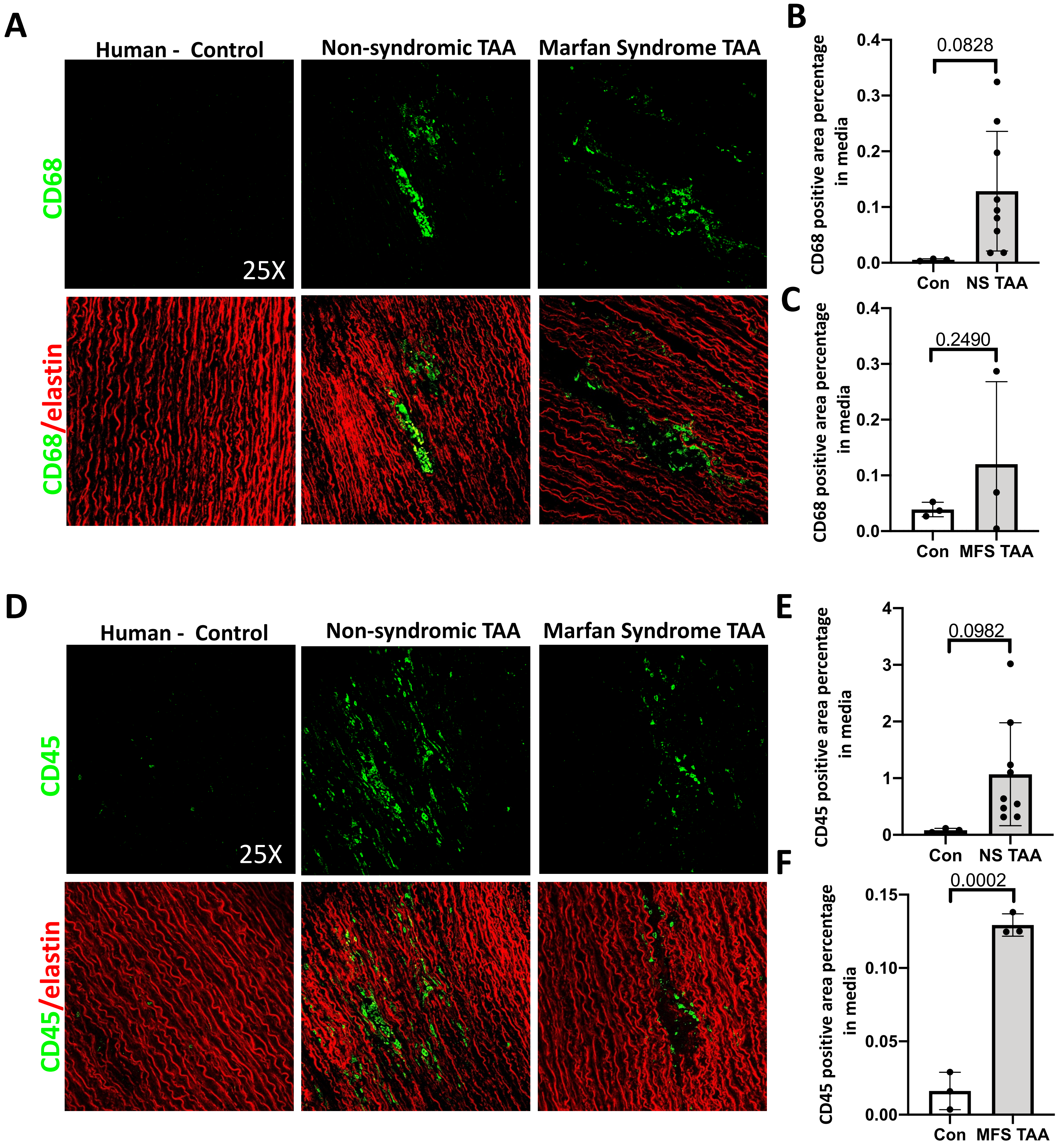
A
Human - Control
Non-syndromic TAA
Marfan Syndrome TAA
CD68
CD68/elastin
25X
B
CD68 positive area percentage in media
0.0828
0.0
0.1
0.2
0.3
0.4
Con
NS TAA
C
CD68 positive area percentage in media
0.2490
Con
MFS TAA
D
Human - Control
Non-syndromic TAA
Marfan Syndrome TAA
CD45
CD45/elastin
25X
E
CD45 positive area percentage in media
0.0982
0
1
2
3
4
Con
NS TAA
F
CD45 positive area percentage in media
0.0002
0.00
0.05
0.10
0.15
Con
MFS TAA

### Table S1

| | ATA (Ascending Thoracic Aorta) | | | |
|---|---|---|---|---|
| | **WT** | **α5/2** | **mgR** | **α5/2 mgR** |
| | **n = 6** | **n = 5** | **n = 5** | **n = 5** |
| **Unloaded dimensions** | | | | |
| Wall Thickness (µm) | 107 ± 1.8 | 112 ± 5.4 | 146 ± 7.8 | 122 ± 7.5 |
| Outer Diameter (µm) | 1129 ± 28 | 1115 ± 29 | 1801 ± 101 | 1467 ± 184 |
| Axial Length (mm) | 2.15 ± 0.14 | 2.20 ± 0.16 | 3.57 ± 0.10 | 2.85 ± 0.23 |
| **Loaded dimensions** | **P = 120** | **P = 120** | **P = 120** | **P = 120** |
| Outer Diameter (µm) | 1766 ± 27.1 | 1732 ± 60.9 | 2667 ± 124.0 | 2095 ± 255.8 |
| Wall Thickness (µm) | 32 ± 1.0 | 32 ± 1.0 | 59 ± 6.5 | 56 ± 4.3 |
| Inner Radius (µm) | 850 ± 14.3 | 834 ± 29.9 | 1275 ± 59.7 | 992 ± 129.5 |
| *in vivo* Axial Stretch ($\lambda_z^{iv}$) | 1.96 ± 0.07 | 2.06 ± 0.10 | 1.61 ± 0.09 | 1.44 ± 0.06 |
| *in vivo* Circumferential Stretch ($\lambda_\vartheta$) | 1.70 ± 0.04 | 1.70 ± 0.03 | 1.58 ± 0.04 | 1.53 ± 0.02 |
| **Systolic Cauchy Stresses (kPa)** | | | | |
| Circumferential, $\sigma_\vartheta$ | 423 ± 18.7 | 415 ± 13.1 | 361 ± 35.5 | 296 ± 52.7 |
| Axial, $\sigma_z$ | 395 ± 24.8 | 348 ± 13.8 | 206 ± 27.6 | 165 ± 24.5 |
| **Systolic Linearized Stiffness (MPa)** | | | | |
| Circumferential, $\mathcal{C}_{\theta\theta\theta\theta}$ | 2.11 ± 0.10 | 2.24 ± 0.10 | 7.13 ± 1.31 | 5.86 ± 1.97 |
| Axial, $\mathcal{C}_{zzzz}$ | 2.06 ± 0.14 | 1.65 ± 0.08 | 1.29 ± 0.12 | 1.11 ± 0.17 |
| **Distensibility (1/MPa)** | 32.76 ± 1.83 | 29.90 ± 1.74 | 8.09 ± 0.93 | 9.96 ± 2.61 |
| **Stored Energy (kPa)** | 129 ± 9.5 | 122 ± 5.9 | 52 ± 7.8 | 39 ± 5.4 |
| **Loaded dimensions** | **P = 80** | **P = 80** | **P = 80** | **P = 80** |
| Outer Diameter (µm) | 1523 ± 19.9 | 1512 ± 46.2 | 2569 ± 130.8 | 2014 ± 260.3 |
| Wall Thickness (µm) | 38 ± 1.0 | 37 ± 1.3 | 61 ± 6.7 | 59 ± 4.5 |
| Inner Radius (µm) | 724 ± 10.7 | 719 ± 22.3 | 1223 ± 63.1 | 948 ± 132.2 |
| *in vivo* Axial Stretch ($\lambda_z^{iv}$) | 1.96 ± 0.07 | 2.06 ± 0.10 | 1.61 ± 0.09 | 1.44 ± 0.06 |
| *in vivo* Circumferential Stretch ($\lambda_\vartheta$) | 1.46 ± 0.04 | 1.47 ± 0.02 | 1.52 ± 0.04 | 1.46 ± 0.02 |
| **Systolic Cauchy Stresses (kPa)** | | | | |
| Circumferential, $\sigma_\vartheta$ | 206 ± 8.4 | 207 ± 6.7 | 222 ± 21.3 | 181 ± 35.1 |
| Axial, $\sigma_z$ | 294 ± 20.3 | 253 ± 10.9 | 157 ± 22.4 | 128 ± 21.9 |
| **Systolic Linearized Stiffness (MPa)** | | | | |
| Circumferential, $\mathcal{C}_{\theta\theta\theta\theta}$ | 0.92 ± 0.04 | 0.94 ± 0.04 | 2.22 ± 0.25 | 1.91 ± 0.54 |
| Axial, $\mathcal{C}_{zzzz}$ | 1.42 ± 0.10 | 1.08 ± 0.05 | 0.91 ± 0.10 | 0.82 ± 0.14 |
| **Stored Energy (kPa)** | 83 ± 6.1 | 80 ± 3.9 | 41 ± 6.0 | 30 ± 5.2 |
| **Energy Dissipation (%)** | 6 ± 0.9 | 6 ± 0.6 | 13 ± 12.6 | 9 ± 8.7 |

### Table S2

| | DTA (Descending Thoracic Aorta) | | | |
|---|---|---|---|---|
| | **WT** | **α5/2** | **mgR** | **α5/2 mgR** |
| | **n = 6** | **n = 6** | **n = 6** | **n = 6** |
| **Unloaded dimensions** | | | | |
| Wall Thickness (µm) | 102 ± 2.2 | 93 ± 1.6 | 97 ± 5.9 | 85 ± 4.2 |
| Outer Diameter (µm) | 800 ± 28 | 806 ± 20 | 836 ± 19 | 789 ± 26 |
| Axial Length (mm) | 3.54 ± 0.29 | 3.74 ± 0.18 | 4.95 ± 0.39 | 4.39 ± 0.35 |
| **Loaded dimensions** | **P = 120** | **P = 120** | **P = 120** | **P = 120** |
| Outer Diameter (µm) | 1215 ± 51.4 | 1296 ± 52.2 | 1277 ± 45.6 | 1208 ± 45.5 |
| Wall Thickness (µm) | 37 ± 2.0 | 32 ± 1.5 | 38 ± 1.5 | 33 ± 1.7 |
| Inner Radius (µm) | 570 ± 26.8 | 616 ± 26.4 | 600 ± 23.0 | 570 ± 21.8 |
| *in vivo* Axial Stretch ($\lambda_z^{iv}$) | 1.63 ± 0.04 | 1.67 ± 0.08 | 1.51 ± 0.03 | 1.52 ± 0.02 |
| *in vivo* Circumferential Stretch ($\lambda_\vartheta$) | 1.69 ± 0.06 | 1.77 ± 0.05 | 1.68 ± 0.04 | 1.67 ± 0.06 |
| **Systolic Cauchy Stresses (kPa)** | | | | |
| Circumferential, $\sigma_\vartheta$ | 249 ± 19.0 | 314 ± 22.4 | 252 ± 14.6 | 275 ± 12.5 |
| Axial, $\sigma_z$ | 221 ± 17.8 | 257 ± 23.3 | 177 ± 4.8 | 203 ± 11.4 |
| **Systolic Linearized Stiffness (MPa)** | | | | |
| Circumferential, $\mathcal{C}_{\theta\theta\theta\theta}$ | 1.71 ± 0.25 | 2.03 ± 0.18 | 2.76 ± 0.16 | 2.68 ± 0.19 |
| Axial, $\mathcal{C}_{zzzz}$ | 1.71 ± 0.18 | 1.69 ± 0.11 | 1.94 ± 0.15 | 2.31 ± 0.21 |
| **Distensibility (1/MPa)** | 23.26 ± 3.75 | 24.78 ± 1.84 | 11.36 ± 0.97 | 13.64 ± 2.60 |
| **Systolic Stored Energy (kPa)** | 62 ± 7.2 | 82 ± 10.1 | 45 ± 3.3 | 48 ± 5.7 |
| **Loaded dimensions** | **P = 80** | **P = 80** | **P = 80** | **P = 80** |
| Outer Diameter (µm) | 1099 ± 50.1 | 1160 ± 48.3 | 1212 ± 42.0 | 1135 ± 43.1 |
| Wall Thickness (µm) | 42 ± 2.1 | 36 ± 1.6 | 40 ± 1.4 | 36 ± 1.7 |
| Inner Radius (µm) | 508 ± 26.2 | 544 ± 24.4 | 566 ± 21.0 | 532 ± 20.3 |
| *in vivo* Axial Stretch ($\lambda_z^{iv}$) | 1.63 ± 0.04 | 1.67 ± 0.08 | 1.51 ± 0.03 | 1.52 ± 0.02 |
| *in vivo* Circumferential Stretch ($\lambda_\vartheta$) | 1.51 ± 0.04 | 1.57 ± 0.04 | 1.59 ± 0.04 | 1.56 ± 0.05 |
| **Systolic Cauchy Stresses (kPa)** | | | | |
| Circumferential, $\sigma_\vartheta$ | 133 ± 11.1 | 164 ± 11.2 | 150 ± 7.9 | 159 ± 4.8 |
| Axial, $\sigma_z$ | 165 ± 11.6 | 188 ± 21.8 | 127 ± 4.5 | 145 ± 7.2 |
| **Systolic Linearized Stiffness (MPa)** | | | | |
| Circumferential, $\mathcal{C}_{\theta\theta\theta\theta}$ | 0.77 ± 0.12 | 0.82 ± 0.07 | 1.24 ± 0.06 | 1.25 ± 0.11 |
| Axial, $\mathcal{C}_{zzzz}$ | 1.10 ± 0.09 | 1.07 ± 0.04 | 1.26 ± 0.10 | 1.62 ± 0.16 |
| **Systolic Stored Energy (kPa)** | 42 ± 4.3 | 55 ± 7.4 | 34 ± 2.2 | 34 ± 2.9 |
| **Energy Dissipation (%)** | 8 ± 0.6 | 8 ± 0.9 | 9 ± 0.6 | 7 ± 0.5 |
